# Adaptive variation in human toll-like receptors is contributed by introgression from both Neandertals and Denisovans

**DOI:** 10.1101/022699

**Authors:** Michael Dannemann, Aida M. Andres, Janet Kelso

## Abstract

Pathogens and the diseases they cause have been among the most important selective forces experienced by humans during their evolutionary history. Although adaptive alleles generally arise by mutation, introgression can also be a valuable source of beneficial alleles. Archaic humans, who lived in Europe and Western Asia for over 200,000 years, were likely well-adapted to the environment and its local pathogens, and it is therefore conceivable that modern humans entering Europe and Western Asia who admixed with them obtained a substantial immune advantage from the introgression of archaic alleles.

Here we document a cluster of three toll-like receptors (*TLR6-TLR1-TLR10)* in modern humans that carries three distinct archaic haplotypes, indicating repeated introgression from archaic humans. Two of these haplotypes are most similar to Neandertal genome, while the third haplotype is most similar to the Denisovan genome. The toll-like receptors are key components of innate immunity and provide an important first line of immune defense against bacteria, fungi and parasites. The unusually high allele frequencies and unexpected levels of population differentiation indicate that there has been local positive selection on multiple haplotypes at this locus. We show that the introgressed alleles have clear functional effects in modern humans; archaic-like alleles underlie differences in the expression of the TLR genes and are associated with reduced microbial resistance and increased allergic disease in large cohorts. This provides strong evidence for recurrent adaptive introgression at the *TLR6-TLR1-TLR10* locus, resulting in differences in disease phenotypes in modern humans.

## Introduction

Modern humans dispersing out of Africa were confronted with new environmental challenges including novel foods, pathogens, and a different climate. They also encountered other human forms, and there is accumulating evidence that interbreeding with Neandertals and Denisovans contributed alleles to the modern human gene pool [1–4]. Two recent studies have provided genome-wide maps of archaic haplotypes in present-day people that are likely to have been introduced by introgression from Neandertals [5, 6]. The discovery of introgression from now extinct human forms into modern humans entering Eurasia raises the possibility that the arriving modern humans may have benefitted from the introduction of alleles that existed in archaic humans who were well-adapted to the environment [1–3, 7]. A few cases of such adaptive introgression have been described, largely involving genes that influence systems interacting directly with the environment. For example, the introgression of a Denisovan haplotype in *EPAS1* has recently been shown to confer altitude adaptation in Tibetans [8], and single locus studies have identified adaptive introgression in genes involved in immunity and metabolism including genes of the Major Histocompatibility Locus (MHC), *SLC16A11*, *OAS1* and *STAT2* [9–12].

Among the top 1% of genes with the highest Neandertal introgression in Eurasians, 12% are immune-related, including three of the ten human toll-like receptors (*TLR-10*, *TLR1* and *TLR-6*) encoded in a cluster on chromosome 4 (Supplementary Table 1). This TLR cluster is particularly interesting because of the extended length of the Neandertal-like haplotype and because of the critical role that the TLRs play innate immunity. The innate immune system provides a first line of defense against pathogens and is involved in the early detection of micro-organisms as well as in the activation of the adaptive immune response [13, 14]. In humans, TLR1, TLR6 and TLR10 occur on the cell surface and are known to detect bacterial, fungal and parasite components including flagellin and glycolipids. They are essential for eliciting the inflammatory and anti-microbial responses as well as for activating an adaptive immune response [15]. In agreement with their crucial role, TLRs have been shown to be highly conserved, although cell-surface TLRs are less strictly constrained than intracellular TLRs [16]. In fact, the region encompassing the *TLR6-TLR1-TLR10* cluster has been shown to have some signatures of recent positive selection in certain non-African populations [16] suggesting that they may be involved in local immune-system adaptation. However, the presence of introgressed archaic haplotypes confounds many of the signatures of recent positive selection [17]. We therefore explored in detail both the evidence for introgression and the evidence for natural selection at the *TLR6-TLR1-TLR10* cluster.

## Results

### Identifying introgressed haplotypes

We identified an extended region encompassing the genes *TLR10*, *TLR1* and *TLR6* on chromosome 4 in both genome-wide maps of Neandertal introgression [5, 6](Figure 1, Supplementary Table 1, Methods). Since the two introgression maps did not agree completely on the length of the introgressed region we used consecutive SNPs with the largest differences in introgression probabilities in the Neandertal introgression map [5] to mark the start and end of a putatively introgressed region that is 143kb in length (hg19, chr4:38760338-38905731, Methods).

**Figure 1:**
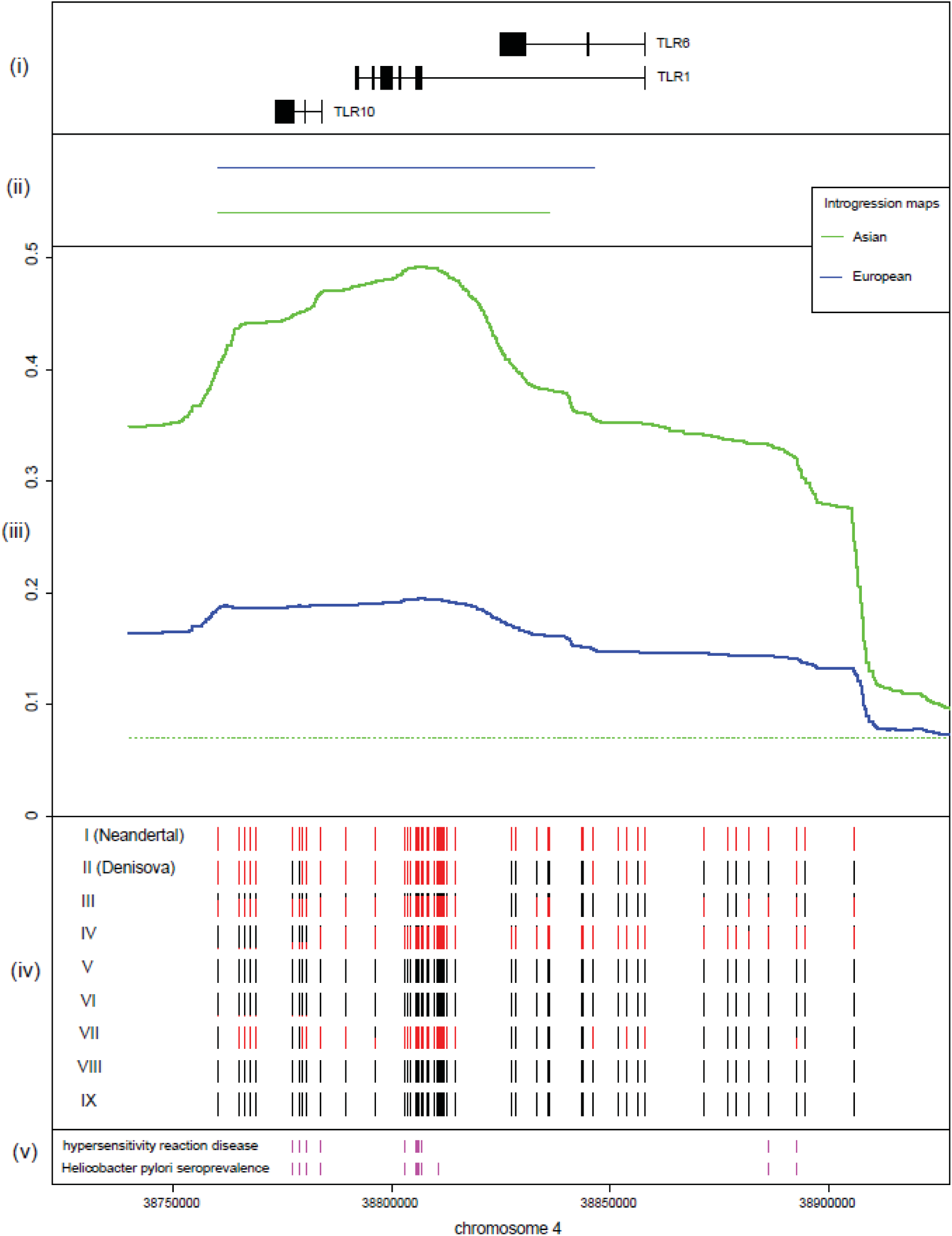
Genomic region on chromosome 4 (*hg19*; 4:38760338-38905731). 
i. The gene structures for *TLR10*, *TLR1* and *TLR6* are displayed.
ii. Predicted Neanderthal haplotypes from the Neanderthal ancestry map by Vernot et al. [6] for Asians (green) and Europeans (blue).
iii. Neanderthal introgression posterior probabilities across polymorphic positions for Asians (blue solid lines) and Europeans (green solid lines) from the Neandertal ancestry map by Sankararaman et al. [5]. The chromosomal average introgression posterior probabilities are given as green and blue dashed lines.
iv. Sharing of the Neanderthal allele across archaic-like SNPs within seven core haplotypes: the Neandertal and Denisova sequences and the seven modern human core haplotypes. The frequencies of Neandertal and Yoruba alleles across the archaic-like SNPs within each core haplotype are colored in red and black, respectively.
v. Significantly associated SNPs with Helicobacter seroprevalence or allergic disease overlapping archaic-like SNPs are displayed [36, 38].

To determine the haplotype structure of this *TLR* region we used the inferred 1,000 Genomes Phase III haplotypes for 2,535 individuals from 26 populations [18]. By clustering haplotypes that differ by fewer than 1/1000 bases we identified seven distinct core haplotypes in modern humans (Methods, Supplementary Figure 1, Supplementary Table 2).

We computed pairwise distances between these seven modern human core haplotypes and the genome sequences of the Neandertal and Denisovan [2, 3] and found that three of the modern human core haplotypes (III, IV and VII) are found almost exclusively in individuals outside Africa and are more similar to the archaic genomes than to any modern human core haplotype (Figure 2, Supplementary Figure 5). In fact, their distances to the archaic sequences are smaller than the distances between any pair of modern human core haplotypes. This suggests a recent common ancestor for the archaic haplotypes and modern human haplotypes III, IV and VII that is younger than the common ancestor of the other pairs of modern human core haplotypes, as expected from archaic introgression. Strikingly, this analysis indicates that there are three distinct archaic-like haplotypes at this locus. Of the three putatively introgressed core haplotypes, III and IV are most similar to the Altai Neanderthal genome (haplotype I). All 49 Neandertal-like alleles that define core haplotype III are shared with core haplotype IV, which has 12 additional archaic-like alleles. Core haplotype VII is most similar to the Denisovan sequence (Figure 2, Supplementary Figure 1, Supplementary Table 5). We find 137 individuals homozygous for core haplotype III and two individuals homozygous for core haplotype IV (IV: HG02388, NA18625, III: Supplementary Table 10), which shows that these haplotypes were not introduced by errors in phasing. We did not identify any individual that was homozygous for core haplotype VII due to its low frequency.

**Figure 2:**
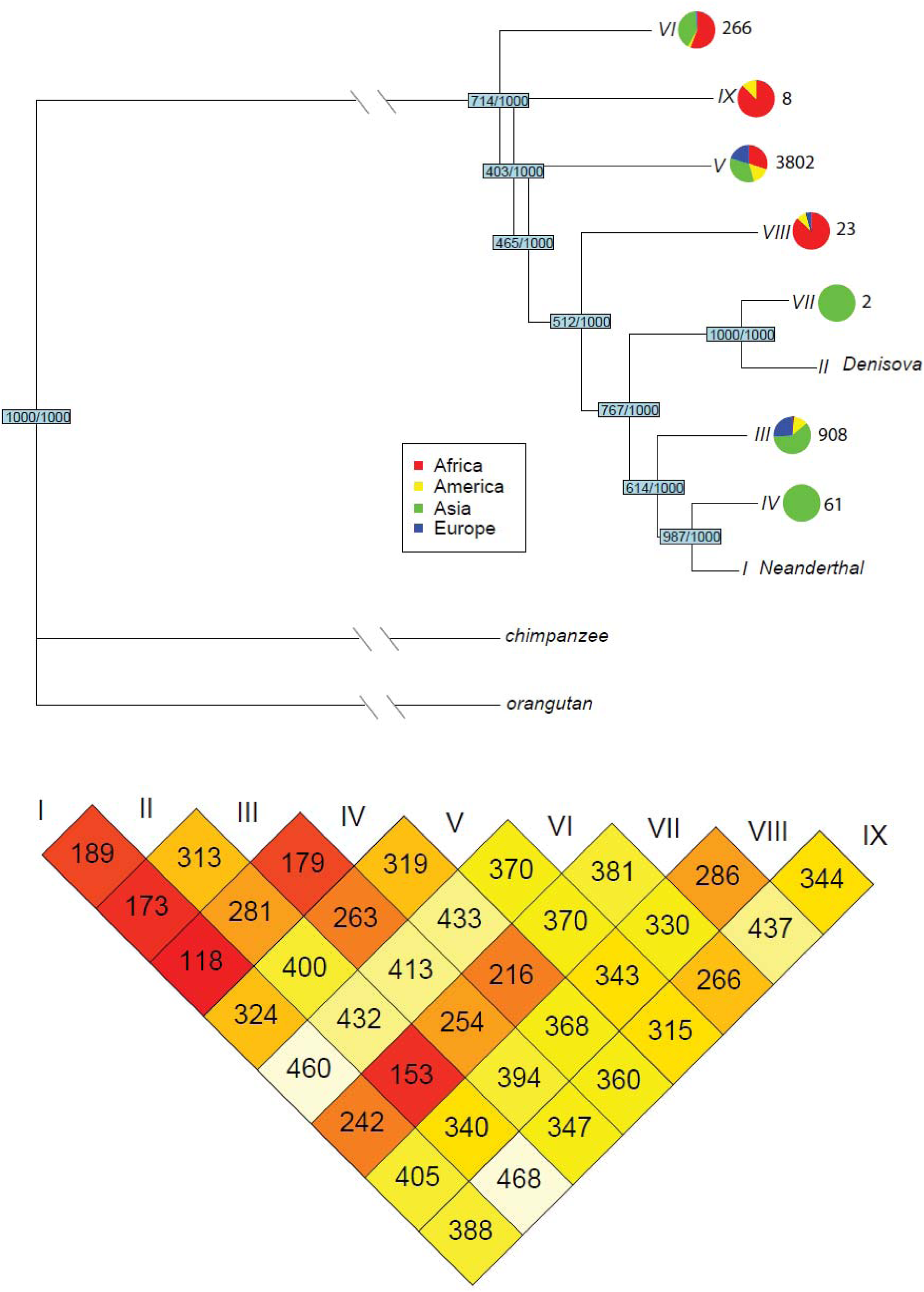
(a) Neighbor joining tree based on the sequences of seven modern human core haplotypes (III-IX), and the Neanderthal (I), Denisova (II), orangutan and chimpanzee genome sequences. Bootstrap values (1000 replicates) for the topology are provided in blue squares at each node. The pie charts show the frequency in the four continental population groups of each modern human core haplotype. Next to each pie chart the frequency of haplotypes among 1,000 genomes individuals assigned to the corresponding core haplotype is displayed. (b) Pair-wise average nucleotide distance between haplotypes in core haplotypes.

### Evidence for introgression

An alternative explanation for the persistence of three archaic-like haplotypes is incomplete lineage sorting (ILS). Both introgression and ILS can result in the presence of ancestral alleles in present-day human genomes, and in a genomic region where the haplotype phylogeny does not match the expected phylogeny [19]. The region described here was identified in two different genome-wide maps of Neandertal introgression using methods that are not sensitive to normal segments of ILS [5, 6]. Nevertheless, we explored a number of key signatures that allow us to distinguish ILS from introgression.

Firstly, given the time since divergence between modern and archaic humans, ILS is expected to be restricted to short genomic regions due to the long-term effects of recombination. We calculated the probability that a haplotype of 143kb in length is not broken by recombination since the common ancestor of modern humans and Neanderthals and/or Denisovans following the approach used by Huerta-Sánchez et al [8]. Using conservative estimates for the age of divergence of the Neandertal and Denisovan (Supplementary Table 12) we show that a 143kb archaic-like haplotype is highly unlikely under ILS (p-value < 2.2×10^−16^, Methods). An unusually low recombination rate in the region is also not a plausible explanation since the recombination rate across the region has been estimated to between 1.5 and 2.4cM/Mb by different methods [20–22] – a rate which is higher than the genome-wide average of 1.3 cM/Mb [23].

Further, evidence in favor of introgression is provided by the presence of archaic-like core haplotypes III, IV and VII almost exclusively in non-Africans. We expect that alleles present due to ILS should be shared across human populations, while Neandertal and Denisova introgressed alleles should be largely absent in Africa. In the 1,000 Genomes Phase III data all three archaic-like haplotypes are largely restricted to populations outside Africa (Figure 2a, 3a, Supplementary Table 5). Core haplotype III is present in all non-African populations, and we also observe it in 2 chromosomes from two Northwest Gambian individuals (excluding the potentially admixed African Americans). Northwest African populations have been shown to have experienced recent gene flow from non-Africans [24] that may have carried archaic introgressed alleles into these populations [25]. This suggests that the presence of this haplotype in these Gambian individuals might be explained by recent back-to-Africa migration. Core haplotype IV is restricted to specific Asian groups, and core haplotype VII is present only in two South Asian individuals (Figure 2a, 3a, Supplementary Table 5)). Haplotypes III and IV are also present among 271 geographically diverse individuals from the Simons Genome Diversity Panel and show a similar geographic distribution (Supplementary Material Figure 3b). The geographic distribution of these archaic-like haplotypes is consistent with recent studies that have inferred two pulses of Neandertal introgression: one into the ancestors of all non-Africans, and a second into the ancestors of present-day Asians [26, 27]. It is also compatible with a low level of Denisovan ancestry in mainland Asia [28]. We note that two early modern humans (dated to ∼7000- 8000 years before present) from Europe (Stuttgart and Loschbour) [29] also carry TLR haplotypes similar to Neandertal core haplotype III while an early modern human from Asia (Ust’-Ishim, ∼45000 years) [30] carries a haplotype that is most similar to core haplotype V – the major non-introgressed haplotype in modern humans (Supplementary Figure 2).

**Figure 3.**
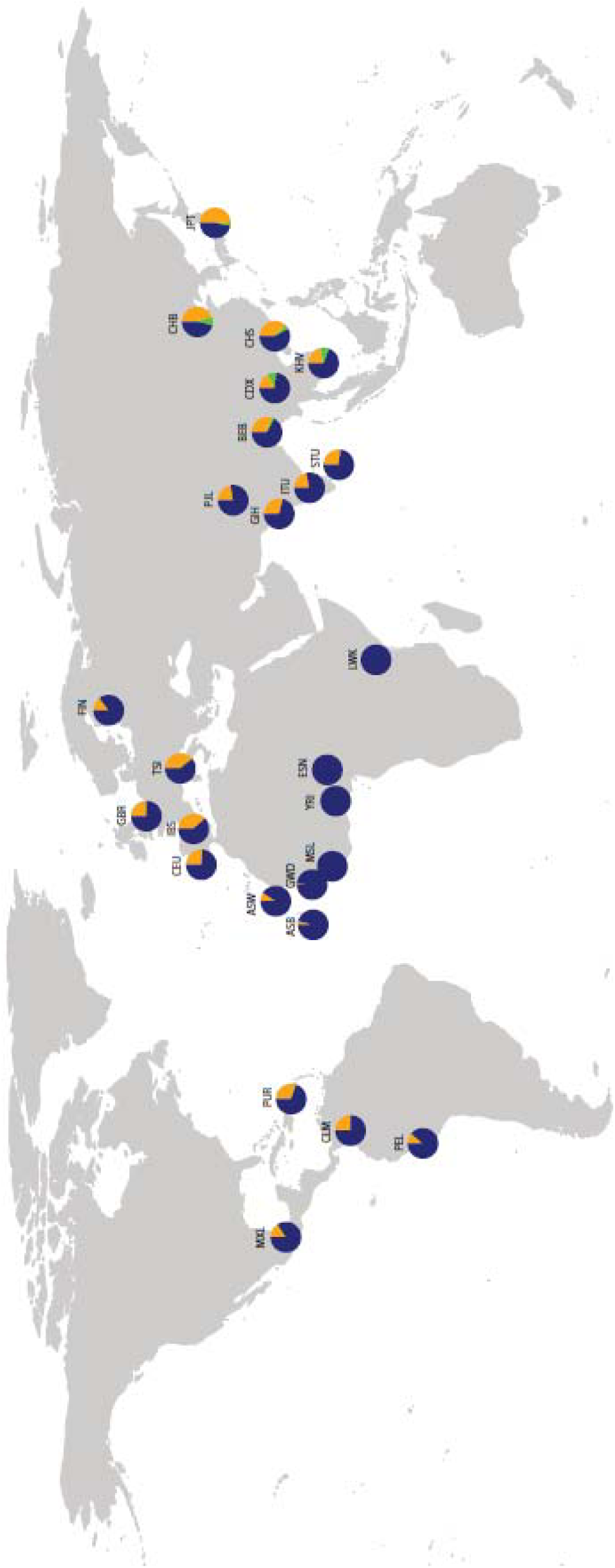
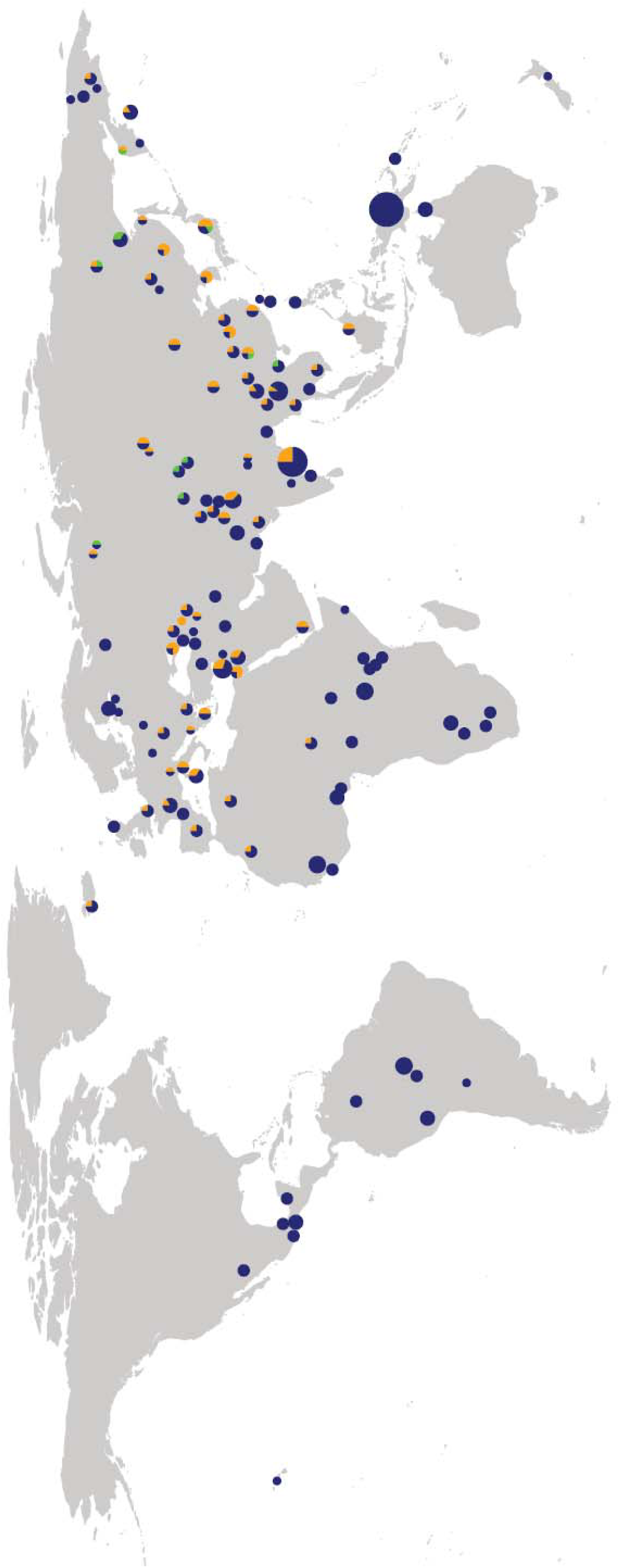
World map showing the frequencies of Neandertal-like core haplotypes in the 1,000 Genomes dataset (a) and the Simons Genome Diversity Panel (b). In (b) the size of each pie is proportional to the number of individuals within a population. Core haplotypes (III=orange, IV=green, non-archaic core haplotypes V,VI,VIII,IX =blue) are colored respectively.

Finally, an expected signature of introgression (but not ILS) is low diversity within the introgressed haplotypes, as they have a recent origin - the introgression event. To assess core haplotype diversity we calculated mean pairwise differences between all haplotypes within each of the seven core haplotype clusters (Supplementary Figure 2, Supplementary Table 11). Consistent with an introgressed origin, the three archaic-like core haplotypes (III, IV and VII) show lower diversity in non-Africans than the most common modern human core haplotype (V) (Supplementary Figure 2, Supplementary Table 11). We also note that core haplotype V has high diversity in Africa (Supplementary Figure 2, Supplementary Table 11). This makes it unlikely that core haplotypes III, IV and VIII were due to ILS and specifically lost in Africans since this would have reduced the overall diversity at this locus.

### Evidence for positive selection in modern humans

The most frequent non-introgressed haplotype (V) is at high frequency in all 1,000 Genomes populations (39-88%), while the other non-introgressed haplotypes (VI, VIII, and IX) are present at less than 1% frequency, and are predominantly found in African individuals (Figure 2, 3, Supplementary Table 5). Among the introgressed haplotypes, Neandertal-like core haplotype III is at intermediate frequency in all non-African populations (11-51%) and Neandertal-like core haplotype IV is present at a frequency of 2-10% in Asians. The Denisovan-like core haplotype VII is present in only two Southern Asian individuals (HG03750, HG0411). The proportion of Neandertal-derived ancestry in non-Africans has been estimated to be between 1.5-2.1%, while Denisovan ancestry in mainland Asia and Native Americans has been estimated at well below 0.5% [3]. The presence of three distinct archaic haplotypes, with two of them at frequencies substantially greater than 2%, is therefore surprising and suggests that the archaic-like alleles may have been advantageous in modern humans.

Recent population-specific selection on the *TLR6-TLR1-TLR10* cluster was reported by Barreiro et al. [16]. However, the presence of introgressed haplotypes, which confound some signatures of selection, was not known at that time. We therefore re-evaluated the signatures of selection at this locus. To determine whether the frequencies of Neandertal-like core haplotypes III and IV (up 10% and 51% in certain Eurasian populations, Supplementary Table 5) are higher than expected from drift alone, we measured allele frequency differentiation between pairs of populations using F_ST_ [31], for all 61 archaic-like SNPs. To assess significance we compared these to the genome-wide empirical distribution of F_ST_ values for putatively introgressed archaic-like SNPs (Methods, Supplementary Table 6). The differentiation between African and non-African populations for the SNPs shared by haplotypes III and IV is unusually high when compared to the empirical F_ST_ distribution of archaic-like SNPs (p-value = 0.01 Africa-Asia and p-value = 0.04-0.05 Africa-Europe, Supplementary Table 6a). Results are similar for core haplotype IV (Supplementary Table 6a). However, because divergence between African and non-African genomes was used as a factor in identifying the region as having high proportion of Neandertal introgression [5], this high population differentiation is not necessarily unexpected. However, we also observe moderate population differentiation between Asians and Europeans for both core haplotype III (p-value=0.08-0.23) and core haplotype IV (p-value=0.11, Supplementary Table 6a), which is not explained by the ascertainment bias.

Perhaps most surprisingly, the frequency of the shared SNPs common to Neandertal-like core haplotypes III and IV also varies significantly between populations even within continents. In Europe, a North-South gradient is apparent with significantly high population differentiation between southern Europeans (Toscani and Iberians) and all other European groups (Finnish, British and CEPH) (p-value <0.05, Supplementary Table 6b). The frequency of the introgressed haplotypes is higher in the Southern populations (Figure 3). In Asia the most Eastern populations (Japanese and Han Chinese) show high differentiation from other Asian populations (p-value<0.05, Supplementary Table 6c). The frequency of the introgressed haplotypes is higher in the Eastern populations. In addition, the 12 SNPs defining Asian-specific core haplotype IV show high population differentiation when Dai and Vietnamese are compared to other Asian populations, when using putatively-introgressed SNPs present exclusively in Asia as the background (p-value<0.05, Supplementary Table 6c). Overall, the frequency of the introgressed haplotypes ranges between 14.8 and 39.3% in Europeans and between 21.7 and 53.5% in Asian populations, although the contribution of each haplotype (III or IV) varies substantially across Asian groups (Figure 3, Supplementary Table 5).

Barreiro et al. [16] previously reported signatures of positive selection tagged by two SNPs: *rs4129009*, a non-synonymous SNP in TLR10 showing signatures of positive selection in East Asians and *rs4833095*, a non-synonymous SNP that reduces TRL1 signaling resulting in a 60% reduction in the activity of NK-kB and shows signatures of positive selection in Europeans. In our data *rs4129009* is predominantly found on Neandertal-like core haplotype III (Supplementary Table 9), and the observed signatures of selection are therefore most likely explained by positive selection on this archaic core haplotype (as shown above). Like Barreiro et al. we observe large population differentiation for *rs4833095*, but in our data this functionally relevant SNP is predominantly found in core haplotype V, which is present in all human populations and therefore unlikely to be introgressed. In fact, the unusually large population differentiation remains when we investigate the frequency of *rs4833095* only within the non-introgressed core haplotype V (Supplementary Table 9). This indicates that the signatures of selection for this SNP are independent of the changes in allele frequency of the introgressed archaic-like core haplotypes (Supplementary Materials)

Together these results provide evidence for multiple independent rounds of positive selection on both introgressed and non-introgressed haplotypes at the *TLR6-TLR1-TLR10* cluster. This region has been proposed to be a hotspot of positive selection in other great apes [32, 33] and the presence of several positively selected alleles in modern human populations is therefore perhaps not surprising. However, the fact that several of these alleles have been acquired through adaptive introgression is quite unusual.

### Functional consequences of the introgressed alleles

There are 42 SNPs that distinguish all archaic-like core haplotypes from the other modern human core haplotypes. Of these, 47% are located upstream of genes, 46% in introns, 2% in 5’UTRs and 2% downstream of genes, 2% in non-coding exons and 1 in 3’UTRs (Methods). None of the archaic-like SNPs modifies the amino acid sequence of any *TLR* gene, but the 143kb encompassing *TLR6-TLR1-TLR10* is rich in regulatory elements, falling in the top 1% of similarly-sized windows for transcription factor binding site density on chromosome 4 [34] (Methods, Supplementary Table 7). In fact, the archaic-like SNPs overlap 49 well-characterized transcription factor binding sites, for 28 different transcription factors, suggesting that any putative functional impact may be regulatory.

To test this hypothesis we obtained expression data for lymphoblastoid cell lines from 421 individuals of European and African origin [35]. All three *TLR* genes show significantly higher expression in individuals carrying archaic-like alleles in core haplotype III than in those carrying the non-introgressed modern human alleles (Supplementary Figure 3, Supplementary Table 4). This is consistent with the observation made in [36] where increased expression of *TLR1* was reported for individuals carrying alleles that we show here are introgressed.

We further assessed the functional relevance of the archaic-like core haplotypes using data from genome-wide association studies. A total of 79 SNPs, of which 13 are archaic-like, are significantly associated with GWAS phenotypes within the 143kb *TLR* region [37]. Of the 37 SNPs significantly associated with *Helicobacter pylori* seroprevalence 13 are archaic-like [36] of the 58 SNPs associated with susceptibility to allergic disease 12 are archaic-like [38] (Figure 1, Supplementary Table 3). Clustering of the seven core haplotypes based on allele sharing with the GWAS SNPs significantly associated with these two phenotypes revealed high sequence differentiation between the three archaic-like core haplotypes and core haplotype V for both phenotypes (Figure 4, Methods). The largest differentiation is between the archaic-like core haplotype III and non-archaic-like core haplotypes, which differ in 23 of the 37 SNPs underlying *Helicobacter pylori* seroprevalence, and in 31 of the 58 SNPs associated with allergic disease (Supplementary Table 3). Interestingly archaic-like alleles are consistently associated with reduced *Helicobacter pylori* seroprevalence [36] and with increased susceptibility to allergic disease [38].

## Discussion

Neandertals lived in Europe and Western Asia for over 200,000 years and were likely well-adapted the environment and local pathogens. It is therefore conceivable that admixture with Neandertals contributed alleles that conferred a substantial immune advantage on modern humans expanding into Europe and Western Asia. The presence of not one, but three distinct introgressed haplotypes, whose sequences are closer to the genomes of two different archaic humans, suggests that maintaining these introgressed alleles is likely to have been advantageous. In fact, the high frequencies that these alleles have reached in several human groups are inconsistent with neutral processes since the admixture between Neandertals and modern humans, around 50,000 years ago [39, 40]. At least two of these introgressed haplotypes appear to have been advantageous in certain modern human populations.

Previous studies highlighted the contribution of archaic human alleles to the adaptive immune system of modern humans [9–12]. We extend this to show that genes involved in innate immunity have also been modified by admixture with archaic humans. These archaic alleles lead to significantly increased expression of *TLR6*, *TLR1* and *TLR10* in white blood cells, and in present-day people are associated with reduced *Helicobacter pylori* seroprevalence and increased susceptibility to allergies [36, 38]. Taken together this suggests that the introgressed alleles may enhance innate immune surveillance and reactivity against certain pathogens, but that this may also have increased hypersensitivity to non-pathogenic allergens, resulting in allergic diseases in present-day people. Given the evidence for selection on multiple human haplotypes and also in other primates, it is possible that exposure to different pathogens may have favored different haplotypes at different times, resulting in higher overall diversity. It has been argued that advantageous genetic diversity can be achieved not only by long-term balancing selection, but also by adaptive introgression from archaic humans [7, 41]. Introgressed haplotypes are a particularly desirable source of diversity because they are both highly divergent, viable, and perhaps even adaptive, in closely related populations [42]. Immune-related loci, which are enriched among targets of balancing selection [41], also likely benefit substantially from the introduction of genetic diversity through means such as introgression [7, 41].

Both adaptive introgression and local positive selection on modern human haplotypes have contributed to the evolution of the *TLR6-TLR1-TLR10* locus in some human populations, affecting both gene expression and protein function. We note, however, that since pathogens evolve quickly it is difficult to pinpoint the precise selective force or forces driving these changes. Further genome-wide screens linking Neandertal haplotypes to modern human phenotypes will provide more insight into how admixture with archaic humans has influenced modern human biology.

## Methods

### Identifying the introgressed region in present day human genomes

To identify potentially introgressed archaic-like haplotypes in present-day human genomes we used the genome-wide Neanderthal introgression maps from Sankararaman et al. and Vernot et al. [5, 6]. The introgression map presented by Sankararaman et al. provides the probability that SNPs at polymorphic positions in modern humans arose on the Neanderthal lineage and introgressed into modern humans. Vernot et al. used the S* statistic [43] to detect putatively introgressed regions in modern humans and compared the candidate regions to the reference Neandertal genome to define introgressed Neandertal blocks in modern humans.

To delineate the introgressed regions of interest we used the per-SNP introgression probabilities from Sankararaman et al. for all Asian and all European individuals. In the region of the three *TLR* genes and an additional region 50kb up- and downstream (chromosome 4: 38723860- 38908438, Figure we computed the difference between the Neandertal probabilities for pairs of neighboring SNPs, relative to the distance between them. We defined the range of our preliminary region of introgression based on the SNPs with the largest difference in Neanderthal probabilities in the introgression maps for both Asians and Europeans. The largest increase was observed at position chr4:38757064 in both the European and Asian maps. The largest decrease in probabilities differed between the European and Asian maps. We used the Asian map (SNP at position chr4: 38907701) as it yielded the longer potentially introgressed region. The maximum difference in the European map is at chr4: 38821916.

We defined archaic-like SNPs as those where the Neandertal or Denisovan differs from the 109 Yoruba individuals in the 1,000 Genomes dataset [18]. We narrowed the potentially introgressed region by selecting the archaic-like SNPs that are within the preliminary region and closest to the borders of this region. The final putatively introgressed region covers 143kb of chromosome 4 (chr4:38760338-38905731; Figure 1) and contains 61 archaic-like SNPs. This region overlaps two haplotypes identified by Vernot et al. - one identified in Europeans and the other in Asians.

### Haplotype network analysis

We constructed a haplotype network based on the haplotypes of all 1,000 genomes individuals (phase III, [18]) and the Altai Neanderthal and Denisovan genome sequences ([44], Supplementary Figure 1). We used all 1,997 SNPs that were observed in more than one chromosome in the 1,000 Genomes dataset [18] and where the Neandertal and the Denisovan were homozygous. We excluded 11 SNPs where either the Neandertal or the Denisovan were heterozygous. We merged the resulting 2,609 unique haplotypes into core haplotypes that differed by <150 nucleotides (i.e.: ∼1/1000 base pairs in the region was allowed to differ between two chromosomes). This resulted in a set of seven modern human core haplotypes (Figure 1, 2 and Supplementary Figure 1, Supplementary Table 2) and two archaic haplotypes (that are simply the Neandertal and Denisovan genome sequences). We generated a consensus sequence for each modern human core haplotype by computing the majority allele at each position.

### Neighbor-joining tree

We computed a Neighbor-Joining tree for the entire 143 kb region based on the nucleotide distance between the sequences of the seven modern human core haplotypes together with the Neanderthal, Denisovan, Chimpanzee and Orangutan sequences using the R package *ape* [45]. We obtained the orthologous sequences for chimpanzee and orangutan by using the liftover software [46]. Branch support was obtained from 1000 bootstraps (Figure 2a).

### Diversity in populations and core haplotypes

We computed the mean pair-wise difference between all haplotype pairs within each core haplotype and for all combinations of populations (Supplementary Figure 2, Supplementary Table 11).

### Recombination

We obtained the average recombination rate from three maps (sex-averaged: deCode 1.5cM/Mb, Marshfield 2.4cM/Mb, Genethon 2.1cM/Mb, [20–22]) for the *TLR* region from UCSC [46]. There is high variation in recombination rates across the region. Two of the maps [22, 47] show a 5-12-fold higher recombination rate than the chromosome average in the region overlapping *TLR10* (Supplementary Figure 4). A third map [48] based on African genomes did not show as extreme an increase in recombination rate (Supplementary Figure 4), which might be partially due to the absence of the archaic-like core haplotypes in individuals with African origin.

### Incomplete lineage sorting

We calculated the probability that a haplotype of 143kb in length is not broken by recombination since the common ancestor of modern humans and Neanderthals and/or Denisovans, following the approach used by Huerta-Sánchez et al [8]. The following parameters were used:

- Recombination rate: *r* = 1.5×10e^-8^. We used the lowest average recombination rate estimate for the test region from the available maps as this yields longer haplotypes and is therefore most conservative.
- Estimates of branch length in years and fossil ages are taken from Prüfer et al. [3] and were tested for mutation rates of both μ=1×10^−9^ and μ = 0.5×10^−9^ per base pair per year.
- Length of the region: 143,000 base pairs
- Generation time: 25 years

### *F*_*ST*_ *analysis*

We computed pairwise F_st_ for all genome-wide SNPs between populations

i. from different continents (Africa, Asia, Europe)
ii. from populations within a continent (Asia and Europe).

F_st_ values were computed using *vcftools* based on the Weir and Cockerham calculation [49].

For (i) we included 100 random unrelated individuals (1,000 Genomes phase III) representing populations from each continent. For (ii) we used 50 unrelated individuals per population. We compared the F_st_ values for two different sets of SNPs

- the 61 archaic-like SNPs that were present in any of the three archaic-like core haplotypes (III, IV, VII) in the region (Supplementary Table 6)
- the four tag SNPs presented in Barreiro et al. [16] (Supplementary Table 9)

Each SNP was compared to the empirical distributions of F_st_ values of:

- all genome-wide SNPs
- all genome-wide archaic-like SNPs
- all genome-wide archaic-like SNPs exclusively found in Asian individuals

We repeated the Fst analyses using all 155 SNPs differentiating the core haplotypes in the region and confirmed the results, solely based on archaic-like SNPs.

### SNPs with evidence for positive selection presented by Barreiro et al

We re-evaluated the signatures for haplotype-defining SNPs reported by Barreiro et al. and their relationship to the core haplotypes we identified. Frequency information for SNPs presented in Barreiro et al. [16] was computed using all individuals of the 1,000 Genomes dataset (phase III) and are shown in Table 9 b-d.

### Expression analysis using a lymphoblastoid cell line data set

We analyzed expression using data from lymphoblastoid cell lines from a subset of 421 Yoruban and European individuals from the 1,000 Genomes cohort [35].

We tested allele-specific expression for the 32 archaic-like SNPs which are shared by the archaic-like core haplotypes (III, IV, VII) and not seen in the other modern-human core haplotypes. Archaic-like SNPs private to core haplotypes IV and VII were not polymorphic in the genomes of individuals of the expression dataset, likely due to their lower frequency and the absence of Asian individuals in this dataset. Of the 32 archaic-like SNPs which are shared by the archaic-like core haplotypes (III, IV, VII) and not seen in the other modern-human core haplotypes 22 were variable in the 1,000 Genomes individuals for whom expression data was available. We obtained mapped reads from http://www.geuvadis.org and filtered for reads with high mapping quality (MQ>30). For each individual we used the remaining reads and evaluated expression of *TLR10*, *TLR1* and *TLR6* by assigning reads with mapping coordinates overlapping a *TLR* genes. We normalized the number of reads per individual using the DESeq package [50] and computed differential expression between the individuals carrying the archaic-like allele and all other individuals using a Mann-Whitney-U test. The FDR values are shown in Supplementary Table 4.

### Transcription factor analysis

We computed the number of transcription factor binding sites obtained from the ENCODE project [34] in 150kb sliding windows with a step size of 10kb across chromosome 4. See Supplementary Materials. We overlapped SNPs which were shared by the archaic-like core haplotypes (III, IV, VII) and different from the other modern human core haplotypes with ENCODE-defined transcription factor binding sites [34] (Supplementary Table 7). We found 49 binding sites for 28 TFs overlapping with these SNPs (Supplementary Table 7).

Using the ontology enrichment software FUNC [51] we tested whether transcription factors with binding sites overlapping archaic-like SNPs show evidence for functional enrichment in the gene ontology [52] compared to other transcription factors (124) tested by the ENCODE consortium.

### Overlap with GWAS phenotype association studies

We identified 79 significantly associated GWAS SNPs within the 143kb *TLR* region using GWASdb [37]. We clustered the seven modern human core haplotypes based on allele sharing with the GWAS SNPs significantly associated with Helicobacter pylori seroprevalence and Susceptiblity to allergic disease using principle component analysis (PCA, Figure 4).

**Figure 4:**
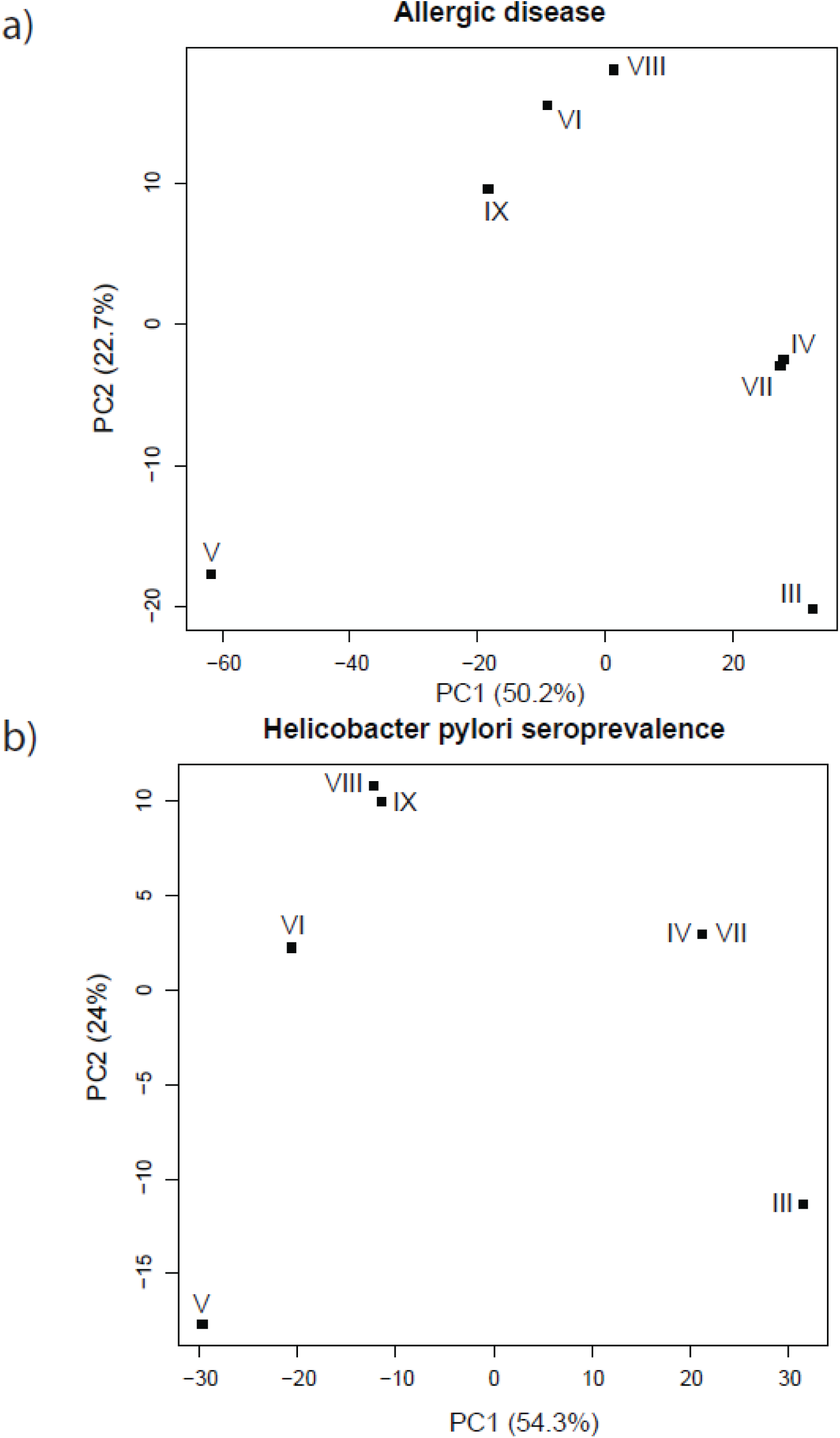
Principal component analysis based on the sequence distance between modern human core haplotypes III-IX for the GWAS SNPs reported to be significantly associated with Helicobacter pylori seroprevalence (a, [36]) and allergic disease (b, [38]).

## Acknowledgments

We thank Cesare de Filippo Felix Key, Svante Päabo, Kay Prüfer, and João Teixeira for helpful discussions. We thank David Reich and Nick Patterson for access to the Simons Genome Diversity Panel data. Funding was provided by the Max Planck Society and the Deutsche Forschungsgemeinschaft SFB1052 “Obesity mechanisms” (project A02) (to J.K).

## Figure and Table Legends

**Table 1**

The rank of average Neanderthal posterior probability (computed using SNPs within genes) for *TLR6-TLR1-TLR10*. A total of 18017 CCDS genes are used in the ranking.

**Table 2:**

(a) 671 variable alleles identified from 2535 individuals in the 1,000 Genomes dataset (b) The majority allele for these 671 variable alleles in each of the nine core haplotypes

**Table 3:**

Column 3: X indicates that the GWAS-identified SNP is an archaic-like SNP

Column 4: X indicates that the SNP is shared by the archaic-like core haplotypes III, IV, VII and differs from the other modern-human core haplotypes II, V, VI, VIII, IX

Column 5: X indicates that the SNP differs between core haplotype III and other modern human core haplotypes II, V, VI, VIII, IX

**Table 4:**

Differential expression FDR value at archaic-like SNP positions and the overlapping gene is displayed.

**Table 5:**

The frequency of seven modern human core haplotypes among 1,000 Genomes population.

**Table 6:**

(a) The table displays the quantile of Fst values for SNPs which differ between the most frequent core haplotype (V) and the archaic-like core haplotypes (III, IV, VII) within the Fst-distribution of

i. genome-wide SNPs,
ii. genome-wide archaic SNPs and
iii. genome-wide-archaic SNPs which are specific to Asians.

Alleles in archaic-like core haplotypes that differ from core haplotype V are highlighted in orange. Quantiles of SNPs within the genome-wide Fst distribution specific to Asians found outside of Asia are shown in grey as these SNPs are not an appropriate background set.

(b) The table displays the quantile of Fst values for SNPs which differ between the most frequent core haplotype (V) and the archaic-like core haplotypes (III, IV, VII) within the Fst-distribution of genome-wide archaic SNPs for the comparisons between European populations.

(c) The table displays the quantile of Fst values for SNPs which differ between the most frequent core haplotype (V) and the archaic-like core haplotypes (III, IV, VII) within the Fst-distribution of genome-wide archaic SNPs for the comparisons between Asian populations.

**Table 7:**

Transcription factor bindings sites overlapping with archaic-like SNPs. Column 1 transcription factor name, columns 2-3: start-end of TF binding site, column 4: position of the overlapping archaic-like SNP.

**Table 8:**

Neanderthal ancestry in Simons Genome Diversity Panel individuals.

For each individual (population identifier in column 1) the number of chromosomes carrying an archaic-like haplotype (columns 2 and 3)

**Table 9:**

Information on the tag SNPs reported by Barreiro et al. [16]

i. Fst values for the pairwise comparisons between 1,000 Genomes populations from Africa, Europe, Asia for the four tag SNPs
ii. The majority allele in each of the nine core haplotypes for the four tag SNPs
iii. Frequency of the human reference allele in each core haplotype for the four tag SNPs
iv. Allele frequency of the human reference allele in modern human populations (1,000 Genomes phase III) for the four tag SNPs

**Table 10:**

List of the 1,000 Genomes individuals that are homozygous for core haplotype III.

**Table 11:**

Nucleotide diversity (in 1×10-4 differences per base pair and the corresponding 95% quantiles) in three *TLR* genes and the entire *TLR* region across core haplotypes.

**Table 12:**

Probability of archaic haplotype block from shared ancestral lineage using different mutation rates. Based on different parameters (age of fossils, recombination rate and estimated branch lengths) we computed the expected length of ILS segments and the probability of our observed haplotypes being a result of ILS.

**Supplementary Figure 1:**
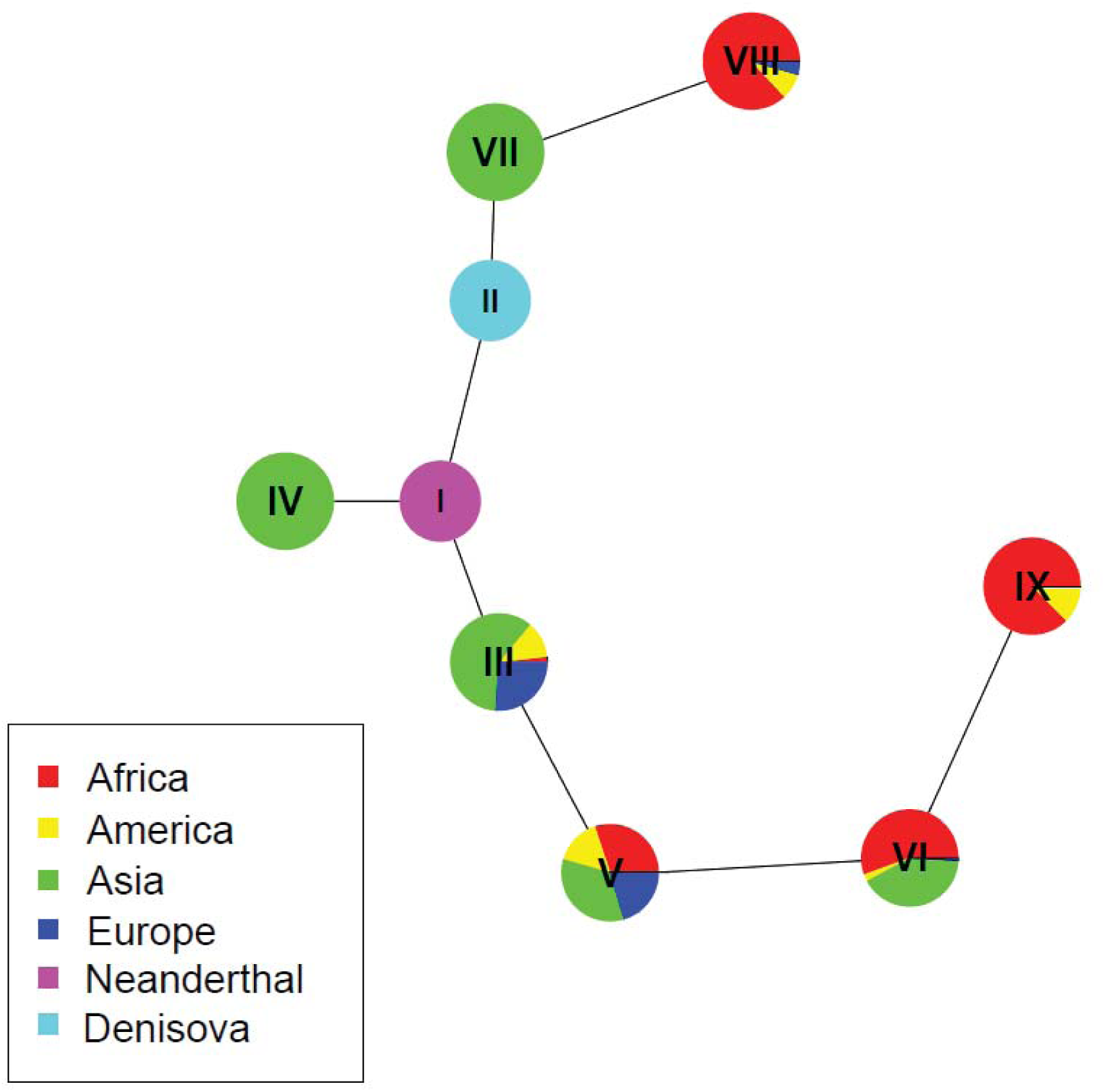
Haplotype network based on the sequence of the nine core haplotypes including the Neandertal (i) and Denisovan (ii) sequences. Colors represent the frequency within each of the 4 super-populations and archaic humans

**Supplementary Figure 2:**
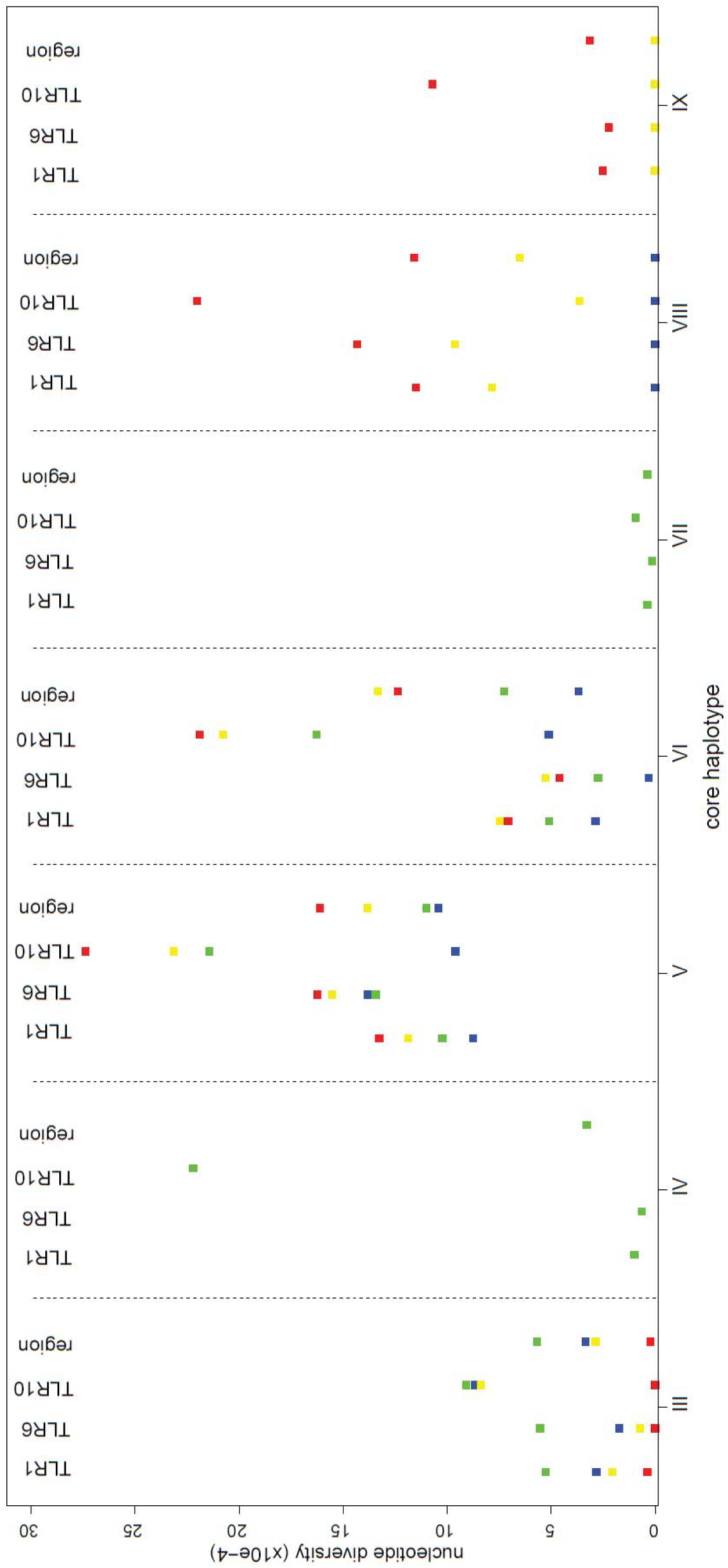
Nucleotide diversity (y-axis) within populations for each core haplotype (x-axis) computed as the mean pair-wise difference between individuals in the respective continent. For each core haplotype and population the nucleotide diversity across the whole region as well as for each *TLR* gene is shown.

**Supplementary Figure 3:**
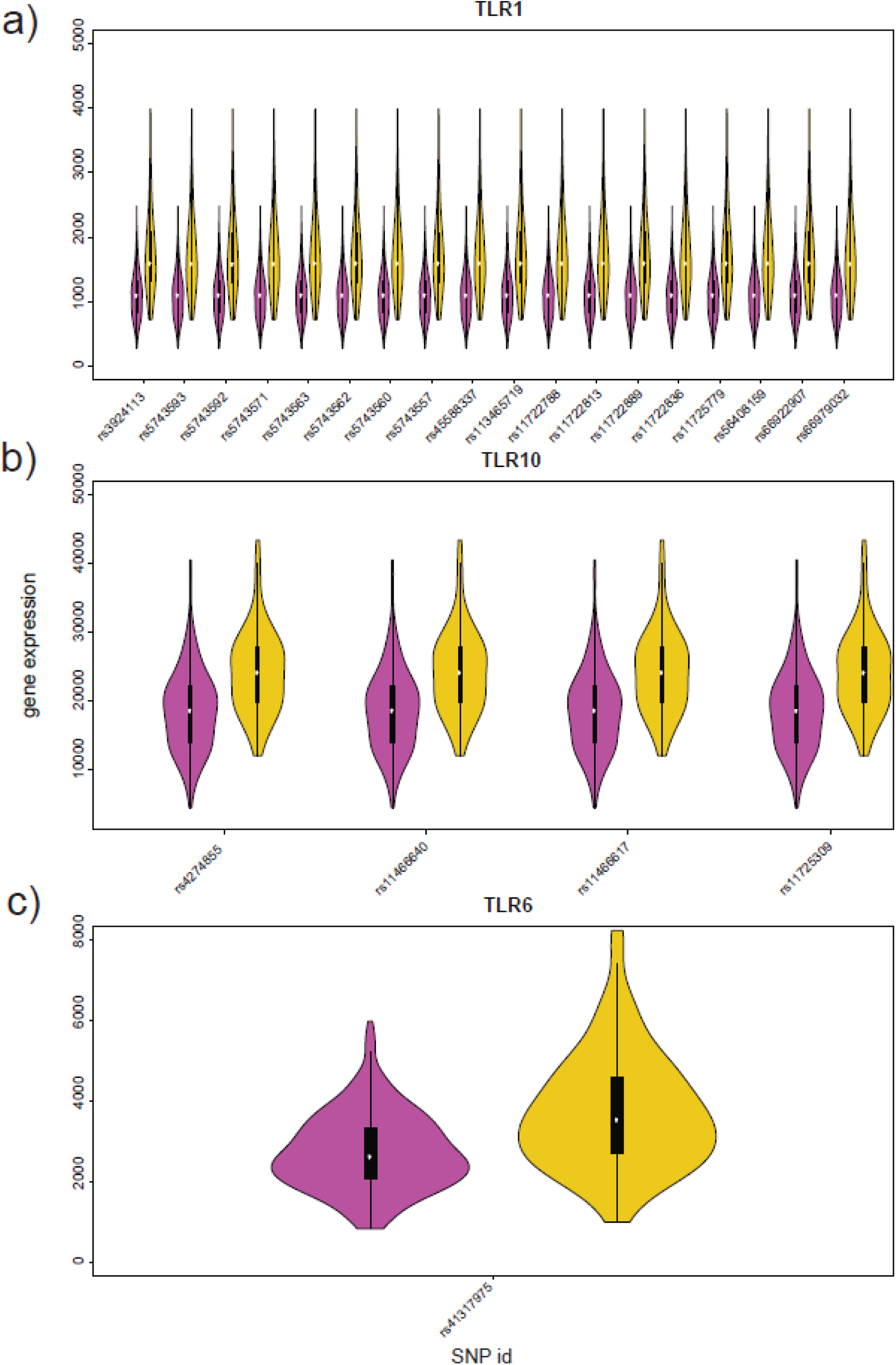
Gene expression in normalised read counts for each of the *TLR* genes (a-c). For all archaic-like SNPs (x-axis SNP rsID) overlapping a *TLR* gene the distribution of expression in individuals carrying an archaic-like allele (magenta) is shown compared to the expression of individuals without the Neanderthal allele (yellow).

**Supplementary Figure 4:**
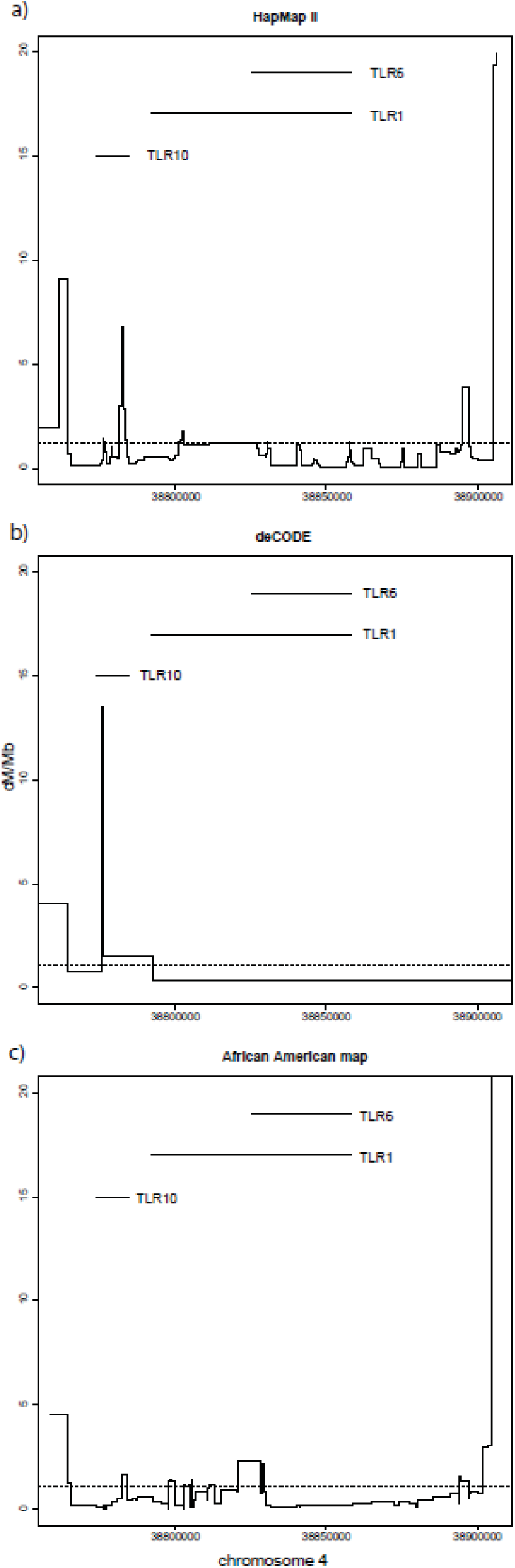
Recombination rate (y-axis) across the *TLR* region (x-axis) for three recombination maps (a-c, [22, 47, 48]) is displayed. The extent of the transcripts of *TLR10*, *TLR6* and *TLR1* are shown on top of the figure. The average recombination rate on chromosome 4 is represented as a dotted line.

**Supplementary Figure 5:**
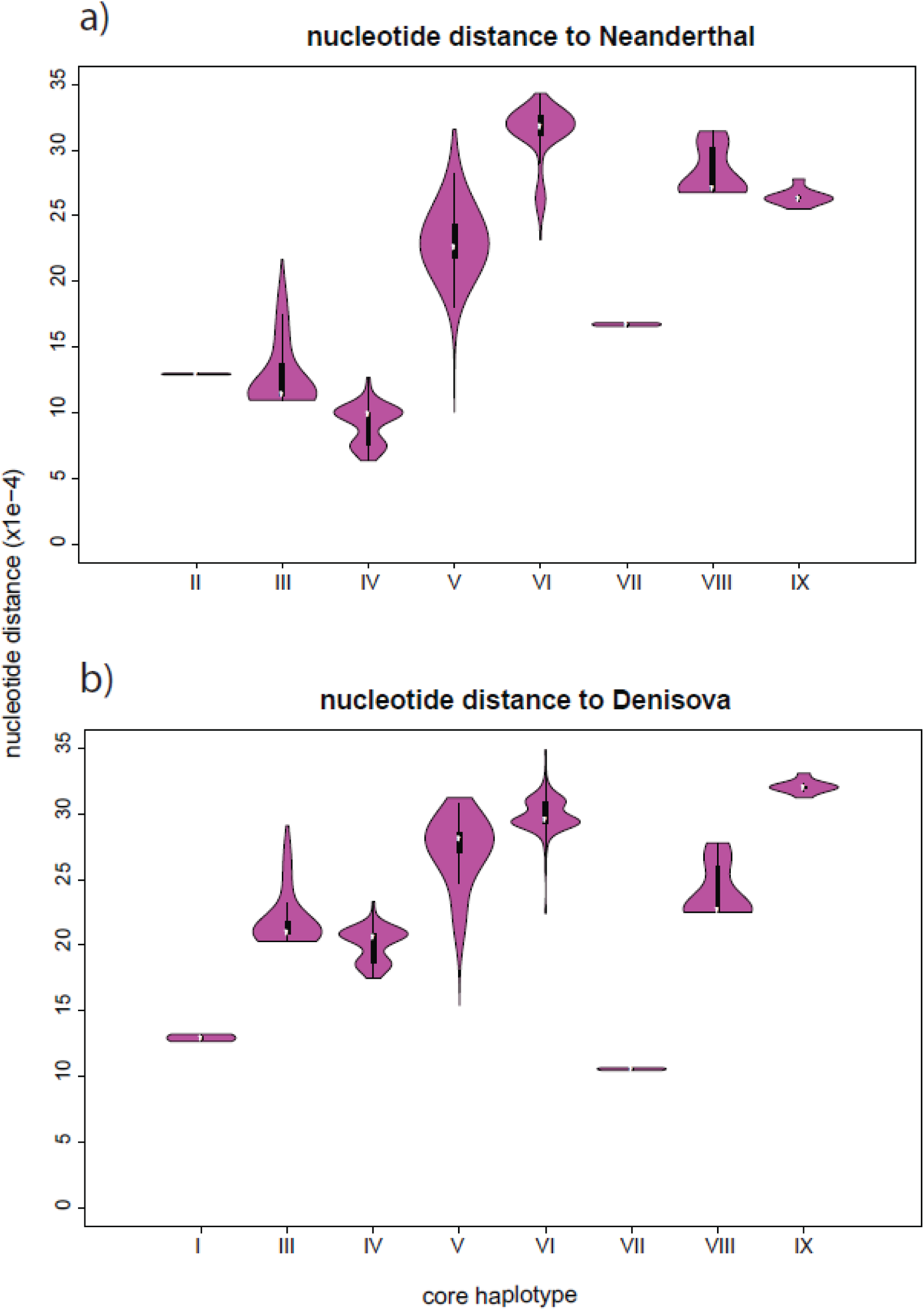
Distributions of the nucleotide distances (y-axis) to the Neanderthal sequence (a) and to the Denisova sequence (b) compared to the remaining eight core haplotypes (x-axis) is shown.

**Supplementary Table 1:** Rank of average introgression scores for *TLR1*, *TLR6* and *TLR10* in Asians and Europeans.

|  | EUR-ASN (combined) | ASN | EUR |
| --- | --- | --- | --- |
| TLR1 | 34 | 40 | 181 |
| TLR6 | 52 | 65 | 279 |
| TLR10 | 39 | 47 | 199 |

**Supplementary Table 2a:**
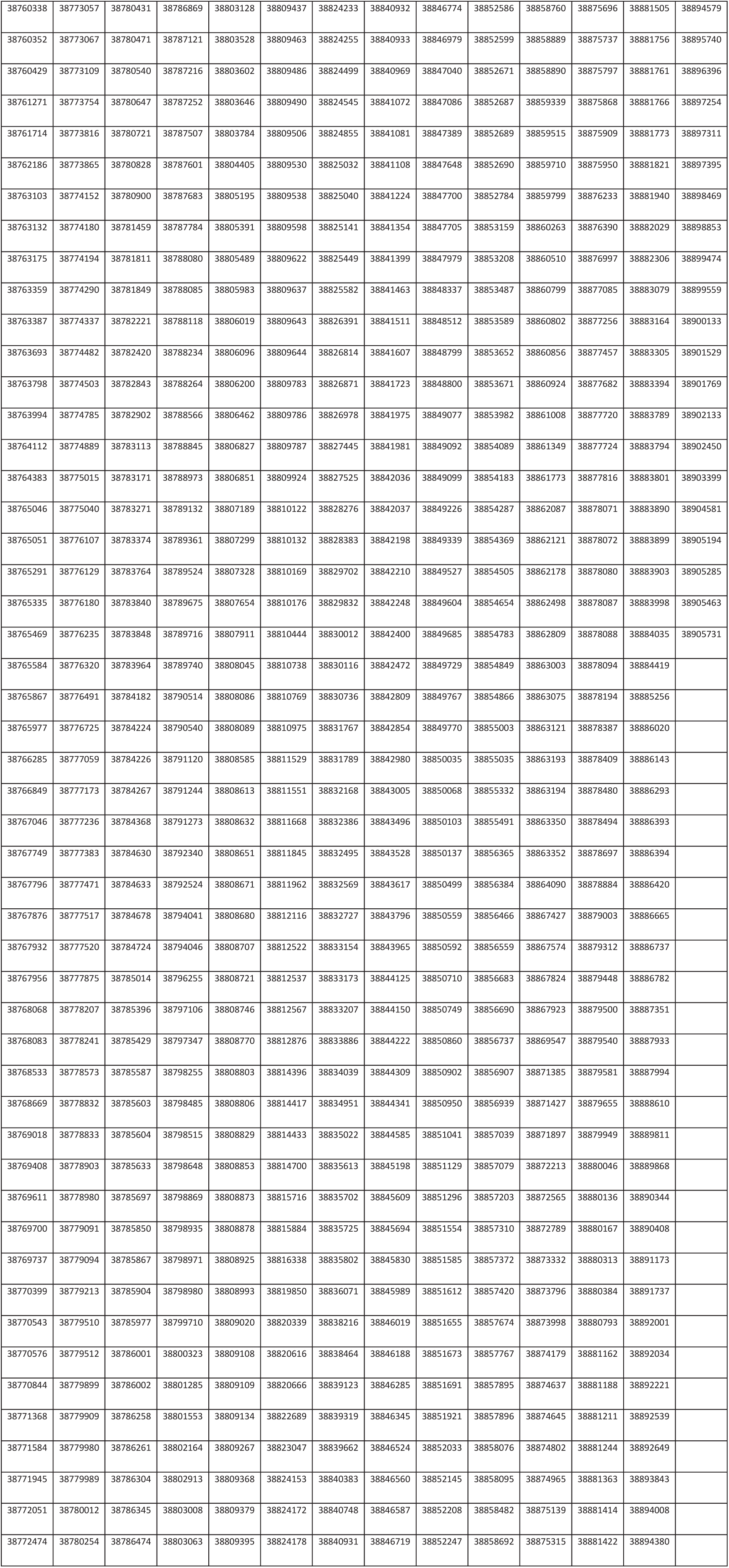
Variable sites in the nine haplotype clusters identified in the *TLR* region.

|  |  |  |  |  |  |  |  |  |  |  |  |  |  |
| --- | --- | --- | --- | --- | --- | --- | --- | --- | --- | --- | --- | --- | --- |
| 38760338 | 38773057 | 38780431 | 38786869 | 38803128 | 38809437 | 38824233 | 38840932 | 38846774 | 38852586 | 38858760 | 38875696 | 38881505 | 38894579 |
| 38760352 | 38773067 | 38780471 | 38787121 | 38803528 | 38809463 | 38824255 | 38840933 | 38846979 | 38852599 | 38858889 | 38875737 | 38881756 | 38895740 |
| 38760429 | 38773109 | 38780540 | 38787216 | 38803602 | 38809486 | 38824499 | 38840969 | 38847040 | 38852671 | 38858890 | 38875797 | 38881761 | 38896396 |
| 38761271 | 38773754 | 38780647 | 38787252 | 38803646 | 38809490 | 38824545 | 38841072 | 38847086 | 38852687 | 38859339 | 38875868 | 38881766 | 38897254 |
| 38761714 | 38773816 | 38780721 | 38787507 | 38803784 | 38809506 | 38824855 | 38841081 | 38847389 | 38852689 | 38859515 | 38875909 | 38881773 | 38897311 |
| 38762186 | 38773865 | 38780828 | 38787601 | 38804405 | 38809530 | 38825032 | 38841108 | 38847648 | 38852690 | 38859710 | 38875950 | 38881821 | 38897395 |
| 38763103 | 38774152 | 38780900 | 38787683 | 38805195 | 38809538 | 38825040 | 38841224 | 38847700 | 38852784 | 38859799 | 38876233 | 38881940 | 38898469 |
| 38763132 | 38774180 | 38781459 | 38787784 | 38805391 | 38809598 | 38825141 | 38841354 | 38847705 | 38853159 | 38860263 | 38876390 | 38882029 | 38898853 |
| 38763175 | 38774194 | 38781811 | 38788080 | 38805489 | 38809622 | 38825449 | 38841399 | 38847979 | 38853208 | 38860510 | 38876997 | 38882306 | 38899474 |
| 38763359 | 38774290 | 38781849 | 38788085 | 38805983 | 38809637 | 38825582 | 38841463 | 38848337 | 38853487 | 38860799 | 38877085 | 38883079 | 38899559 |
| 38763387 | 38774337 | 38782221 | 38788118 | 38806019 | 38809643 | 38826391 | 38841511 | 38848512 | 38853589 | 38860802 | 38877256 | 38883164 | 38900133 |
| 38763693 | 38774482 | 38782420 | 38788234 | 38806096 | 38809644 | 38826814 | 38841607 | 38848799 | 38853652 | 38860856 | 38877457 | 38883305 | 38901529 |
| 38763798 | 38774503 | 38782843 | 38788264 | 38806200 | 38809783 | 38826871 | 38841723 | 38848800 | 38853671 | 38860924 | 38877682 | 38883394 | 38901769 |
| 38763994 | 38774785 | 38782902 | 38788566 | 38806462 | 38809786 | 38826978 | 38841975 | 38849077 | 38853982 | 38861008 | 38877720 | 38883789 | 38902133 |
| 38764112 | 38774889 | 38783113 | 38788845 | 38806827 | 38809787 | 38827445 | 38841981 | 38849092 | 38854089 | 38861349 | 38877724 | 38883794 | 38902450 |
| 38764383 | 38775015 | 38783171 | 38788973 | 38806851 | 38809924 | 38827525 | 38842036 | 38849099 | 38854183 | 38861773 | 38877816 | 38883801 | 38903399 |
| 38765046 | 38775040 | 38783271 | 38789132 | 38807189 | 38810122 | 38828276 | 38842037 | 38849226 | 38854287 | 38862087 | 38878071 | 38883890 | 38904581 |
| 38765051 | 38776107 | 38783374 | 38789361 | 38807299 | 38810132 | 38828383 | 38842198 | 38849339 | 38854369 | 38862121 | 38878072 | 38883899 | 38905194 |
| 38765291 | 38776129 | 38783764 | 38789524 | 38807328 | 38810169 | 38829702 | 38842210 | 38849527 | 38854505 | 38862178 | 38878080 | 38883903 | 38905285 |
| 38765335 | 38776180 | 38783840 | 38789675 | 38807654 | 38810176 | 38829832 | 38842248 | 38849604 | 38854654 | 38862498 | 38878087 | 38883998 | 38905463 |
| 38765469 | 38776235 | 38783848 | 38789716 | 38807911 | 38810444 | 38830012 | 38842400 | 38849685 | 38854783 | 38862809 | 38878088 | 38884035 | 38905731 |
| 38765584 | 38776320 | 38783964 | 38789740 | 38808045 | 38810738 | 38830116 | 38842472 | 38849729 | 38854849 | 38863003 | 38878094 | 38884419 |  |
| 38765867 | 38776491 | 38784182 | 38790514 | 38808086 | 38810769 | 38830736 | 38842809 | 38849767 | 38854866 | 38863075 | 38878194 | 38885256 |  |
| 38765977 | 38776725 | 38784224 | 38790540 | 38808089 | 38810975 | 38831767 | 38842854 | 38849770 | 38855003 | 38863121 | 38878387 | 38886020 |  |
| 38766285 | 38777059 | 38784226 | 38791120 | 38808585 | 38811529 | 38831789 | 38842980 | 38850035 | 38855035 | 38863193 | 38878409 | 38886143 |  |
| 38766849 | 38777173 | 38784267 | 38791244 | 38808613 | 38811551 | 38832168 | 38843005 | 38850068 | 38855332 | 38863194 | 38878480 | 38886293 |  |
| 38767046 | 38777236 | 38784368 | 38791273 | 38808632 | 38811668 | 38832386 | 38843496 | 38850103 | 38855491 | 38863350 | 38878494 | 38886393 |  |
| 38767749 | 38777383 | 38784630 | 38792340 | 38808651 | 38811845 | 38832495 | 38843528 | 38850137 | 38856365 | 38863352 | 38878697 | 38886394 |  |
| 38767796 | 38777471 | 38784633 | 38792524 | 38808671 | 38811962 | 38832569 | 38843617 | 38850499 | 38856384 | 38864090 | 38878884 | 38886420 |  |
| 38767876 | 38777517 | 38784678 | 38794041 | 38808680 | 38812116 | 38832727 | 38843796 | 38850559 | 38856466 | 38867427 | 38879003 | 38886665 |  |
| 38767932 | 38777520 | 38784724 | 38794046 | 38808707 | 38812522 | 38833154 | 38843965 | 38850592 | 38856559 | 38867574 | 38879312 | 38886737 |  |
| 38767956 | 38777875 | 38785014 | 38796255 | 38808721 | 38812537 | 38833173 | 38844125 | 38850710 | 38856683 | 38867824 | 38879448 | 38886782 |  |
| 38768068 | 38778207 | 38785396 | 38797106 | 38808746 | 38812567 | 38833207 | 38844150 | 38850749 | 38856690 | 38867923 | 38879500 | 38887351 |  |
| 38768083 | 38778241 | 38785429 | 38797347 | 38808770 | 38812876 | 38833886 | 38844222 | 38850860 | 38856737 | 38869547 | 38879540 | 38887933 |  |
| 38768533 | 38778573 | 38785587 | 38798255 | 38808803 | 38814396 | 38834039 | 38844309 | 38850902 | 38856907 | 38871385 | 38879581 | 38887994 |  |

|  |  |  |  |  |  |  |  |  |  |  |  |  |
| --- | --- | --- | --- | --- | --- | --- | --- | --- | --- | --- | --- | --- |
| 38768669 | 38778832 | 38785603 | 38798485 | 38808806 | 38814417 | 38834951 | 38844341 | 38850950 | 38856939 | 38871427 | 38879655 | 38888610 |
| 38769018 | 38778833 | 38785604 | 38798515 | 38808829 | 38814433 | 38835022 | 38844585 | 38851041 | 38857039 | 38871897 | 38879949 | 38889811 |
| 38769408 | 38778903 | 38785633 | 38798648 | 38808853 | 38814700 | 38835613 | 38845198 | 38851129 | 38857079 | 38872213 | 38880046 | 38889868 |
| 38769611 | 38778980 | 38785697 | 38798869 | 38808873 | 38815716 | 38835702 | 38845609 | 38851296 | 38857203 | 38872565 | 38880136 | 38890344 |
| 38769700 | 38779091 | 38785850 | 38798935 | 38808878 | 38815884 | 38835725 | 38845694 | 38851554 | 38857310 | 38872789 | 38880167 | 38890408 |
| 38769737 | 38779094 | 38785867 | 38798971 | 38808925 | 38816338 | 38835802 | 38845830 | 38851585 | 38857372 | 38873332 | 38880313 | 38891173 |
| 38770399 | 38779213 | 38785904 | 38798980 | 38808993 | 38819850 | 38836071 | 38845989 | 38851612 | 38857420 | 38873796 | 38880384 | 38891737 |
| 38770543 | 38779510 | 38785977 | 38799710 | 38809020 | 38820339 | 38838216 | 38846019 | 38851655 | 38857674 | 38873998 | 38880793 | 38892001 |
| 38770576 | 38779512 | 38786001 | 38800323 | 38809108 | 38820616 | 38838464 | 38846188 | 38851673 | 38857767 | 38874179 | 38881162 | 38892034 |
| 38770844 | 38779899 | 38786002 | 38801285 | 38809109 | 38820666 | 38839123 | 38846285 | 38851691 | 38857895 | 38874637 | 38881188 | 38892221 |
| 38771368 | 38779909 | 38786258 | 38801553 | 38809134 | 38822689 | 38839319 | 38846345 | 38851921 | 38857896 | 38874645 | 38881211 | 38892539 |
| 38771584 | 38779980 | 38786261 | 38802164 | 38809267 | 38823047 | 38839662 | 38846524 | 38852033 | 38858076 | 38874802 | 38881244 | 38892649 |
| 38771945 | 38779989 | 38786304 | 38802913 | 38809368 | 38824153 | 38840383 | 38846560 | 38852145 | 38858095 | 38874965 | 38881363 | 38893843 |
| 38772051 | 38780012 | 38786345 | 38803008 | 38809379 | 38824172 | 38840748 | 38846587 | 38852208 | 38858482 | 38875139 | 38881414 | 38894008 |
| 38772474 | 38780254 | 38786474 | 38803063 | 38809395 | 38824178 | 38840931 | 38846719 | 38852247 | 38858692 | 38875315 | 38881422 | 38894380 |

**Supplementary Table 2b:**
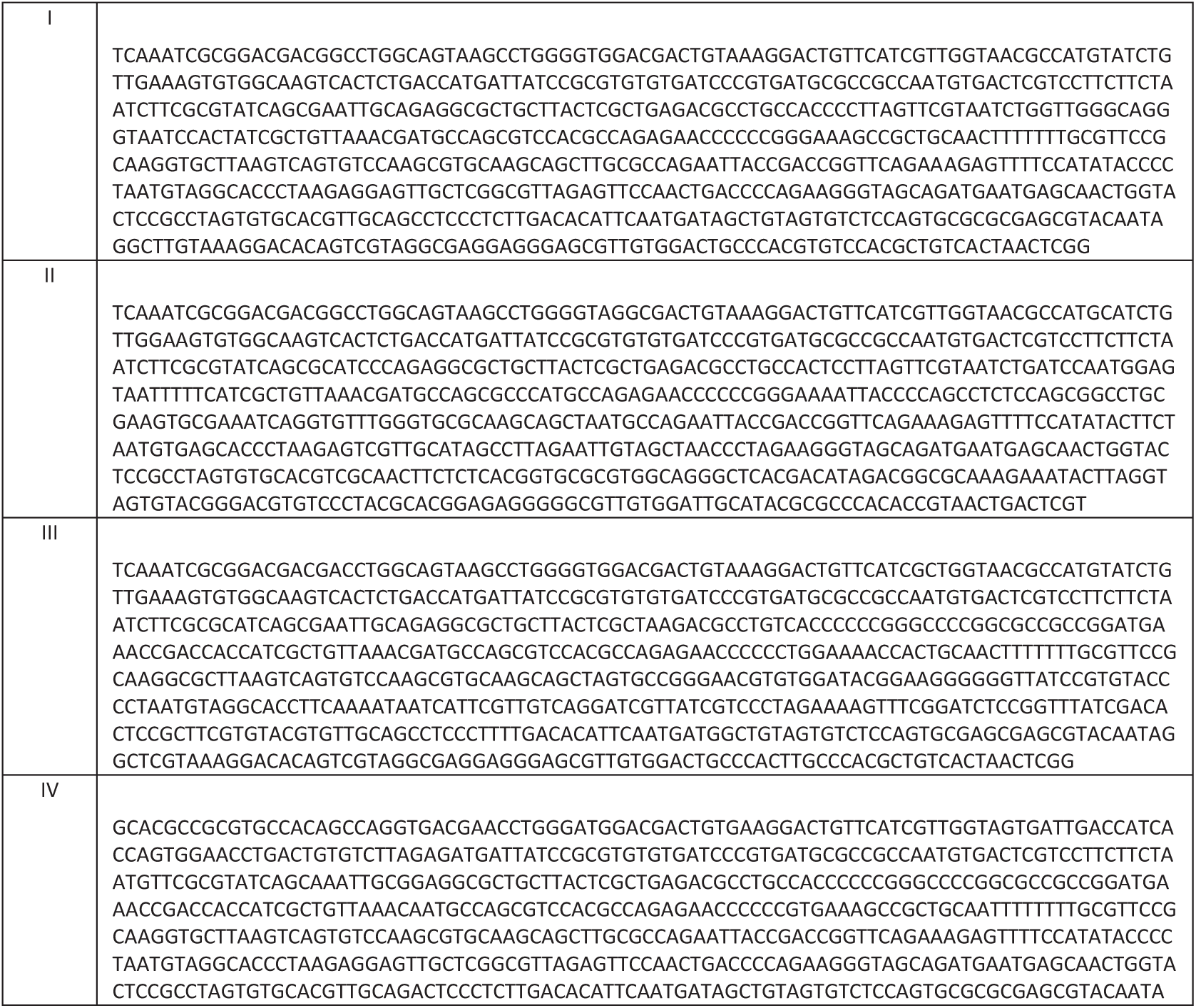
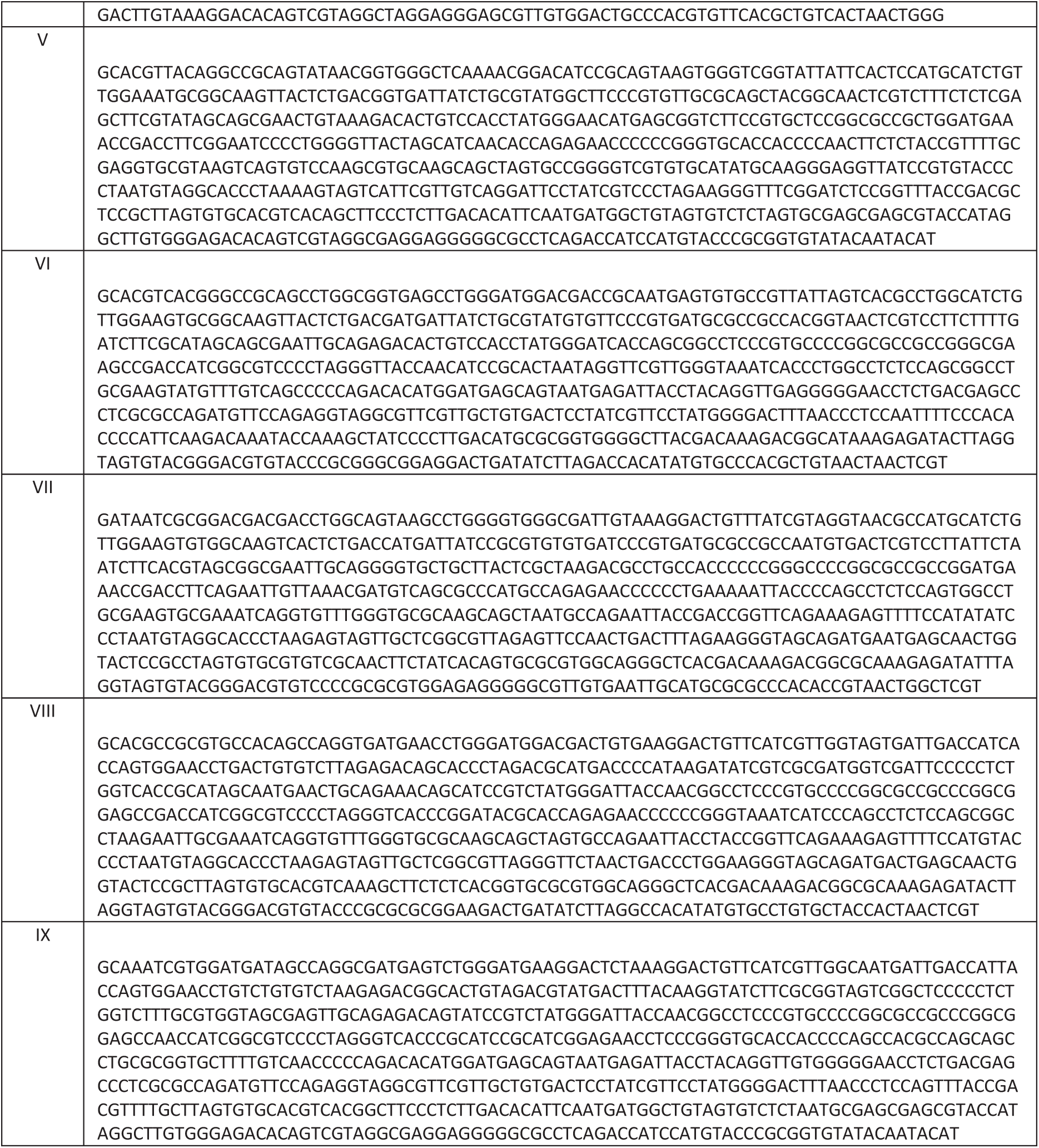
Sequences corresponding to the variable sites in the nine core haplotypes identified in the *TLR* region

**Supplementary Table 3:**
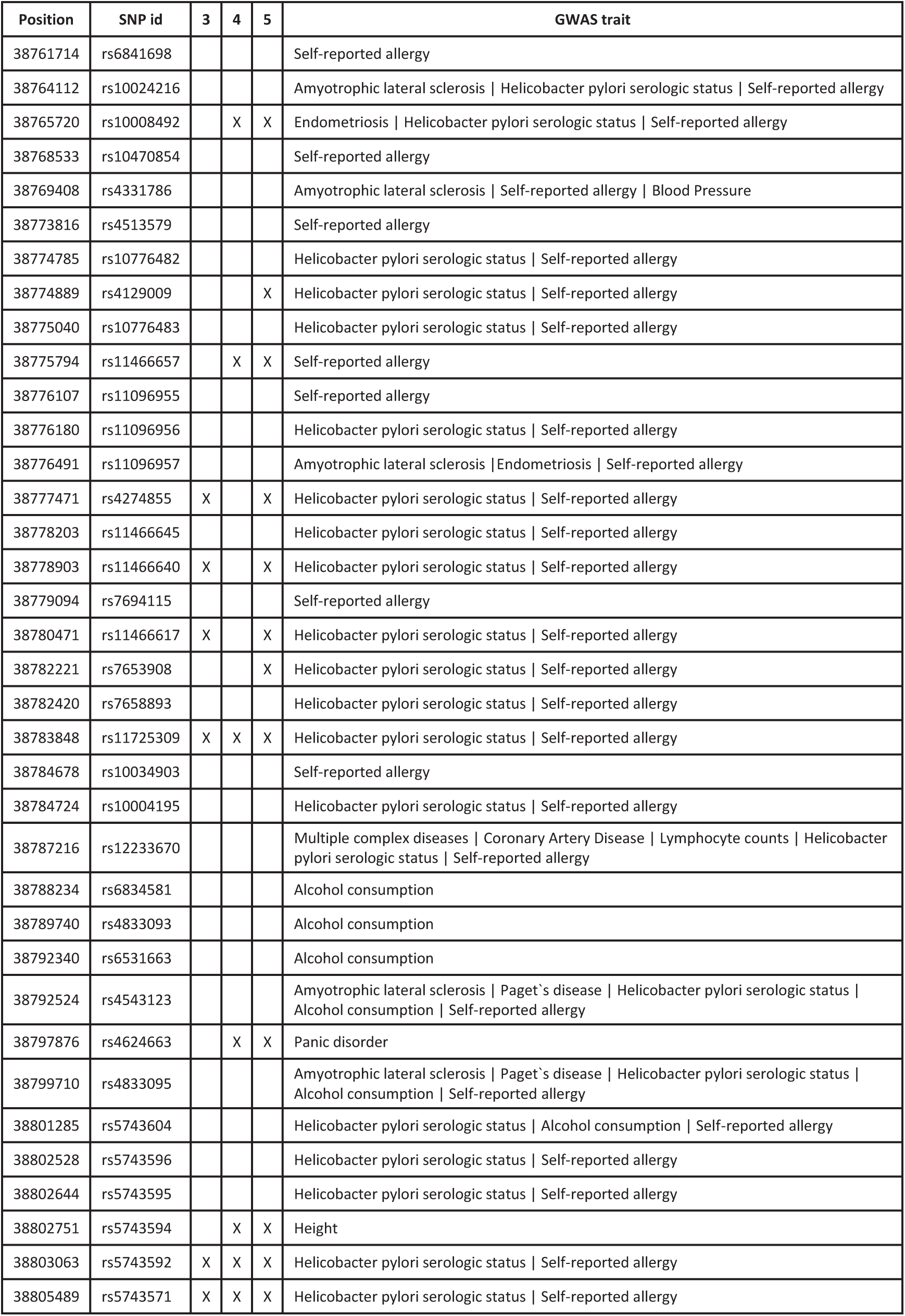
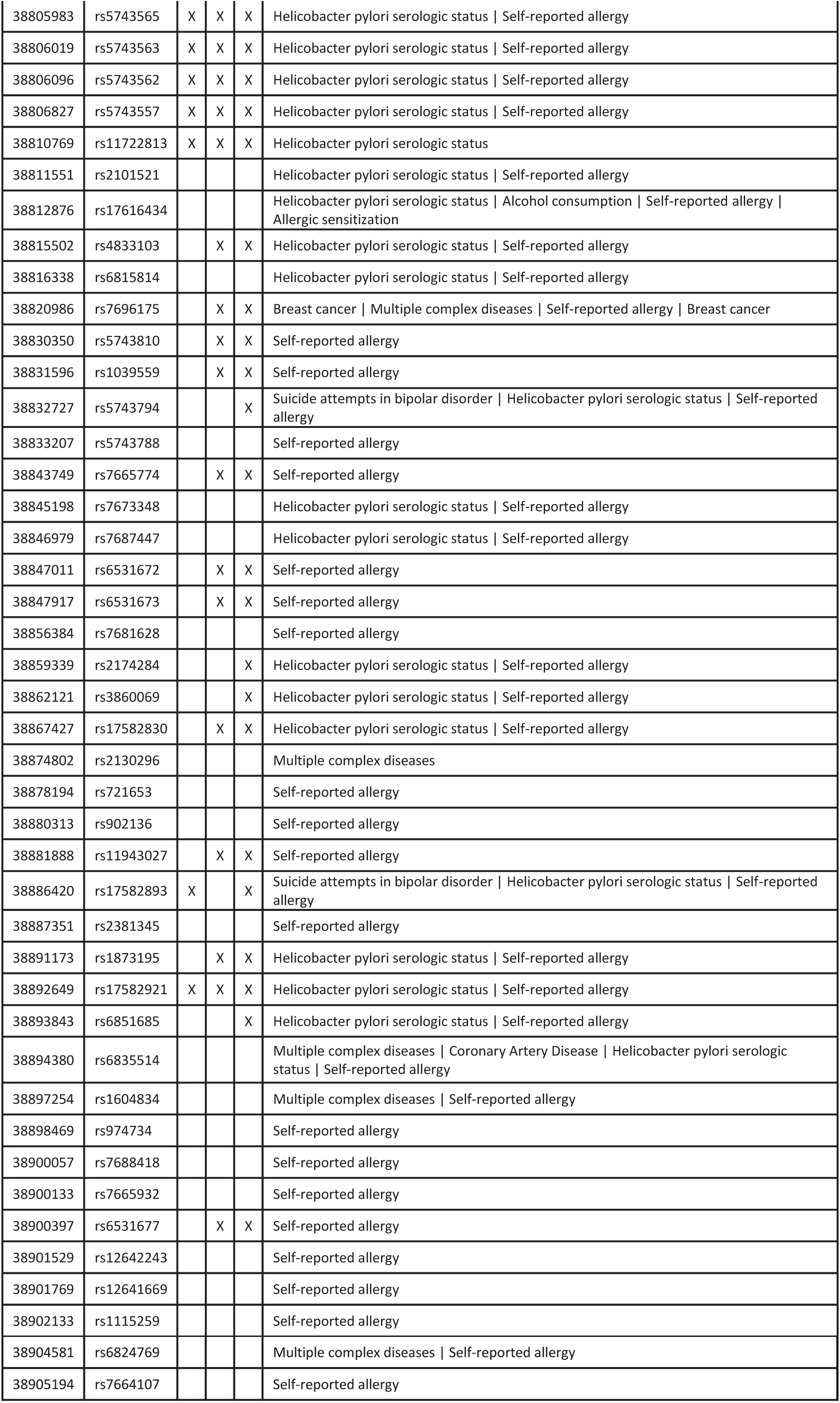
Significant GWAS SNPs and associated GWAS traits in the TLR region and their overlap with archaic-like core haplotypes.

**Supplementary Table 4:** Differential expression of *TLR* genes between individuals with and without the archaic-like allele listed in column 1

| Position | SNP id | Gene | FDR |
| --- | --- | --- | --- |
| 38796255 | rs3924113 | TLR1 | 2.6e-23 |
| 38802913 | rs5743593 | TLR1 | 5.0e-23 |
| 38803063 | rs5743592 | TLR1 | 1.1e-22 |
| 38805489 | rs5743571 | TLR1 | 5.0e-23 |
| 38806019 | rs5743563 | TLR1 | 2.6e-23 |
| 38806096 | rs5743562 | TLR1 | 2.6e-23 |
| 38806200 | rs5743560 | TLR1 | 2.6e-23 |
| 38806827 | rs5743557 | TLR1 | 2.6e-23 |
| 38807328 | rs45588337 | TLR1 | 2.6e-23 |
| 38808045 | rs113465719 | TLR1 | 5.4e-23 |
| 38810738 | rs11722788 | TLR1 | 5.4e-23 |
| 38810769 | rs11722813 | TLR1 | 5.4e-23 |
| 38810975 | rs11722889 | TLR1 | 5.4e-23 |
| 38811529 | rs11722836 | TLR1 | 5.4e-23 |
| 38811668 | rs11725779 | TLR1 | 5.4e-23 |
| 38812116 | rs56408159 | TLR1 | 5.4e-23 |
| 38812537 | rs66922907 | TLR1 | 5.4e-23 |
| 38814700 | rs66979032 | TLR1 | 2.0e-22 |
| 38777471 | rs4274855 | TLR10 | 3.2e-15 |
| 38778903 | rs11466640 | TLR10 | 3.2e-15 |
| 38780471 | rs11466617 | TLR10 | 3.2e-15 |
| 38783848 | rs11725309 | TLR10 | 4.7e-15 |
| 38825443 | rs73236628 | TLR6 | 5.6e-12 |

**Supplementary Table 5:** The frequency of the seven modern human core haplotypes in 1000 genomes populations

|  | III | IV | V | VI | VII | VIII | IX |
| --- | --- | --- | --- | --- | --- | --- | --- |
| CHB | 44.6 | 9 | 40.4 | 6 | 0 | 0 | 0 |
| JPT | 50.9 | 2.5 | 39.3 | 7.4 | 0 | 0 | 0 |
| CHS | 37.2 | 4.7 | 45.9 | 12.2 | 0 | 0 | 0 |
| CDX | 17.4 | 9.4 | 60.4 | 12.8 | 0 | 0 | 0 |
| KHV | 21.1 | 9.9 | 57.9 | 11.2 | 0 | 0 | 0 |
| GIH | 28.9 | 0 | 69 | 2.1 | 0 | 0 | 0 |
| PJL | 22.5 | 0 | 71.3 | 6.2 | 0 | 0 | 0 |
| BEB | 30.7 | 2.4 | 63 | 3.9 | 0 | 0 | 0 |
| STU | 27 | 0 | 67.6 | 4.7 | 0.7 | 0 | 0 |
| ITU | 21.7 | 0 | 73.2 | 4.3 | 0.7 | 0 | 0 |
| CEU | 26.4 | 0 | 73.6 | 0 | 0 | 0 | 0 |
| TSI | 39.3 | 0 | 60 | 0.7 | 0 | 0 | 0 |
| FIN | 14.8 | 0 | 85.2 | 0 | 0 | 0 | 0 |
| GBR | 24.8 | 0 | 74.4 | 0.8 | 0 | 0 | 0 |
| IBS | 38.3 | 0 | 61 | 0 | 0 | 0.6 | 0 |
| YRI | 0 | 0 | 85.7 | 12.7 | 0 | 1.6 | 0 |
| LWK | 0 | 0 | 73.1 | 26.9 | 0 | 0 | 0 |
| GWD | 1.5 | 0 | 83 | 10.4 | 0 | 1.5 | 3.7 |
| MSL | 0 | 0 | 76.1 | 17.4 | 0 | 4.6 | 1.8 |
| ESN | 0 | 0 | 83.6 | 12.1 | 0 | 4.3 | 0 |
| ASW | 9.2 | 0 | 65.3 | 24.5 | 0 | 1 | 0 |
| ACB | 4.1 | 0 | 77.7 | 14.9 | 0 | 3.3 | 0 |
| MXL | 15.9 | 0 | 80.5 | 1.2 | 0 | 2.4 | 0 |
| PUR | 30.3 | 0 | 66.9 | 2.8 | 0 | 0 | 0 |
| CLM | 25.2 | 0 | 74 | 0.8 | 0 | 0 | 0 |
| PEL | 11.2 | 0 | 87.8 | 0 | 0 | 0 | 1 |

**Supplementary Table 6a:** Quantiles (1-P-values) for F_st_ values for pair-wise comparisons between Africans, Asians and Europeans.

| chr | Position | core haplotype | | | | genome-wide $F_{ST}$ | | | genome-wide archaic SNPs | | | Asian-specific archaic SNPs | |
| --- | --- | --- | --- | --- | --- | --- | --- | --- | --- | --- | --- | --- | --- |
|  |  | V | III | IV | VII | ASN EUR | ASN AFR | EUR AFR | ASN EUR | ASN AFR | EUR AFR | ASN EUR | ASN AFR |
| 4 | 38760338 | G | T | G | G | 0.751 | 0.973 | 0.939 | 0.658 | 0.992 | 0.969 | - | - |
| 4 | 38765046 | A | G | A | G | 0.798 | 0.974 | 0.931 | 0.739 | 0.992 | 0.960 | - | - |
| 4 | 38766285 | G | A | G | A | 0.798 | 0.974 | 0.931 | 0.739 | 0.992 | 0.960 | - | - |
| 4 | 38767749 | G | A | G | A | 0.810 | 0.975 | 0.931 | 0.76 | 0.992 | 0.960 | - | - |
| 4 | 38769018 | A | G | A | G | 0.810 | 0.975 | 0.931 | 0.76 | 0.992 | 0.960 | - | - |
| 4 | 38777471 | C | T | C | C | 0.836 | 0.978 | 0.931 | 0.806 | 0.994 | 0.960 | - | - |
| 4 | 38778903 | G | A | G | G | 0.836 | 0.978 | 0.931 | 0.806 | 0.994 | 0.960 | - | - |
| 4 | 38779512 | C | T | C | T | 0.836 | 0.978 | 0.931 | 0.806 | 0.994 | 0.960 | - | - |
| 4 | 38780471 | T | C | T | C | 0.836 | 0.978 | 0.931 | 0.806 | 0.994 | 0.960 | - | - |
| 4 | 38783848 | T | C | C | C | 0.897 | 0.983 | 0.925 | 0.899 | 0.996 | 0.955 | - | - |
| 4 | 38789675 | G | A | A | A | 0.897 | 0.982 | 0.917 | 0.899 | 0.995 | 0.947 | - | - |
| 4 | 38796255 | G | T | T | G | 0.890 | 0.980 | 0.919 | 0.890 | 0.995 | 0.948 | - | - |
| 4 | 38802913 | A | G | G | G | 0.899 | 0.985 | 0.931 | 0.901 | 0.997 | 0.960 | - | - |
| 4 | 38803063 | A | G | G | G | 0.908 | 0.987 | 0.934 | 0.913 | 0.997 | 0.963 | - | - |
| 4 | 38803646 | T | C | C | C | 0.899 | 0.985 | 0.931 | 0.901 | 0.997 | 0.960 | - | - |
| 4 | 38803784 | C | T | T | T | 0.899 | 0.985 | 0.931 | 0.901 | 0.997 | 0.960 | - | - |
| 4 | 38804405 | C | T | T | T | 0.899 | 0.985 | 0.931 | 0.901 | 0.997 | 0.960 | - | - |
| 4 | 38805489 | C | T | T | T | 0.899 | 0.985 | 0.931 | 0.901 | 0.997 | 0.960 | - | - |
| 4 | 38805983 | T | C | C | C | 0.904 | 0.986 | 0.931 | 0.908 | 0.997 | 0.960 | - | - |
| 4 | 38806019 | A | G | G | G | 0.904 | 0.986 | 0.931 | 0.908 | 0.997 | 0.960 | - | - |
| 4 | 38806096 | T | C | C | C | 0.899 | 0.985 | 0.931 | 0.901 | 0.997 | 0.960 | - | - |
| 4 | 38806200 | G | T | T | T | 0.899 | 0.985 | 0.931 | 0.901 | 0.997 | 0.960 | - | - |
| 4 | 38806827 | G | A | A | A | 0.899 | 0.985 | 0.931 | 0.901 | 0.997 | 0.960 | - | - |
| 4 | 38806851 | A | G | G | G | 0.899 | 0.985 | 0.931 | 0.901 | 0.997 | 0.960 | - | - |
| 4 | 38807328 | A | G | G | G | 0.899 | 0.985 | 0.931 | 0.901 | 0.997 | 0.960 | - | - |
| 4 | 38808045 | A | T | T | T | 0.905 | 0.985 | 0.928 | 0.909 | 0.997 | 0.957 | - | - |
| 4 | 38808585 | G | C | C | C | 0.905 | 0.985 | 0.928 | 0.909 | 0.997 | 0.957 | - | - |
| 4 | 38808613 | G | A | A | A | 0.905 | 0.984 | 0.92 | 0.909 | 0.996 | 0.950 | - | - |
| 4 | 38809924 | G | A | A | A | 0.911 | 0.985 | 0.925 | 0.917 | 0.997 | 0.955 | - | - |
| 4 | 38810444 | C | T | T | T | 0.905 | 0.985 | 0.928 | 0.909 | 0.997 | 0.957 | - | - |
| 4 | 38810738 | C | G | G | G | 0.905 | 0.985 | 0.928 | 0.909 | 0.997 | 0.957 | - | - |
| 4 | 38810769 | C | T | T | T | 0.905 | 0.985 | 0.928 | 0.909 | 0.997 | 0.957 | - | - |
| 4 | 38810975 | C | T | T | T | 0.905 | 0.985 | 0.928 | 0.909 | 0.997 | 0.957 | - | - |
| 4 | 38811529 | T | A | A | A | 0.905 | 0.985 | 0.928 | 0.909 | 0.997 | 0.957 | - | - |
| 4 | 38811668 | G | A | A | A | 0.905 | 0.985 | 0.928 | 0.909 | 0.997 | 0.957 | - | - |
| 4 | 38811845 | G | C | C | C | 0.905 | 0.985 | 0.928 | 0.909 | 0.997 | 0.957 | - | - |
| 4 | 38812116 | T | A | A | A | 0.905 | 0.985 | 0.928 | 0.909 | 0.997 | 0.957 | - | - |
| 4 | 38812537 | A | G | G | G | 0.905 | 0.985 | 0.928 | 0.909 | 0.997 | 0.957 | - | - |
| 4 | 38814700 | A | G | G | G | 0.914 | 0.986 | 0.928 | 0.922 | 0.997 | 0.957 | - | - |
| 4 | 38827445 | A | A | G | A | 0.799 | 0.735 | - | 0.741 | 0.708 | - | 0.890 | 0.889 |
| 4 | 38828383 | A | A | G | A | 0.799 | 0.735 | - | 0.741 | 0.708 | - | 0.890 | 0.889 |
| 4 | 38833173 | A | T | T | A | 0.865 | 0.981 | 0.931 | 0.854 | 0.995 | 0.960 | - | - |
| 4 | 38835702 | T | C | C | T | 0.865 | 0.981 | 0.931 | 0.854 | 0.995 | 0.960 | - | - |
| 4 | 38836071 | G | A | A | G | 0.865 | 0.981 | 0.931 | 0.854 | 0.995 | 0.960 | - | - |
| 4 | 38843617 | A | A | T | A | 0.799 | 0.735 | - | 0.741 | 0.708 | - | 0.890 | 0.889 |
| 4 | 38843965 | T | T | C | T | 0.799 | 0.735 | - | 0.741 | 0.708 | - | 0.890 | 0.889 |
| 4 | 38846188 | T | T | G | G | 0.799 | 0.735 | - | 0.741 | 0.708 | - | 0.890 | 0.889 |
| 4 | 38851921 | T | T | G | T | 0.799 | 0.735 | - | 0.741 | 0.708 | - | 0.890 | 0.889 |
| 4 | 38853671 | G | G | A | A | 0.799 | 0.735 | - | 0.741 | 0.708 | - | 0.890 | 0.889 |
| 4 | 38856365 | T | T | C | T | 0.799 | 0.735 | - | 0.741 | 0.708 | - | 0.890 | 0.889 |
| 4 | 38857896 | C | C | A | A | 0.799 | 0.735 | - | 0.741 | 0.708 | - | 0.890 | 0.889 |
| 4 | 38871385 | T | C | C | T | 0.849 | 0.980 | 0.934 | 0.826 | 0.994 | 0.963 | - | - |
| 4 | 38876997 | G | G | A | G | 0.799 | 0.735 | - | 0.741 | 0.708 | - | 0.890 | 0.889 |
| 4 | 38879003 | A | A | C | A | 0.799 | 0.735 | - | 0.741 | 0.708 | - | 0.890 | 0.889 |
| 4 | 38881761 | G | A | A | G | 0.845 | 0.978 | 0.928 | 0.821 | 0.994 | 0.957 | - | - |
| 4 | 38881766 | G | A | A | G | 0.845 | 0.978 | 0.928 | 0.821 | 0.994 | 0.957 | - | - |
| 4 | 38881773 | A | G | G | A | 0.827 | 0.976 | 0.928 | 0.788 | 0.993 | 0.957 | - | - |
| 4 | 38886420 | G | A | A | G | 0.815 | 0.977 | 0.934 | 0.768 | 0.993 | 0.963 | - | - |
| 4 | 38892649 | T | C | C | C | 0.815 | 0.977 | 0.934 | 0.768 | 0.993 | 0.963 | - | - |
| 4 | 38894579 | C | C | T | C | 0.799 | 0.735 | - | 0.741 | 0.708 | - | 0.890 | 0.889 |
| 4 | 38905731 | T | G | G | T | 0.815 | 0.977 | 0.934 | 0.768 | 0.993 | 0.963 | - | - |

**Supplementary Table 6b:** Quantiles (1-P-values) for F_st_ values for pair-wise comparisons between European populations.

|  |  | core haplotype |  |  |  | population comparisons |  |  |  |  |  |  |  |  |  |
| --- | --- | --- | --- | --- | --- | --- | --- | --- | --- | --- | --- | --- | --- | --- | --- |
| chr | position | V | III | IV | VII | TSI<br>CEU | FIN<br>CEU | FIN<br>TSI | GBR<br>CEU | GBR<br>TSI | GBR<br>FIN | IBS<br>CEU | IBS<br>TSI | IBS<br>FIN | IBS<br>GBR |
| 4 | 38760338 | G | T | G | G | 0.923 | 0.629 | 0.974 | 0.471 | 0.692 | 0.877 | 0.966 | 0.015 | 0.986 | 0.764 |
| 4 | 38765046 | A | G | A | G | 0.96 | 0.58 | 0.984 | 0.2 | 0.919 | 0.712 | 0.773 | 0.405 | 0.941 | 0.66 |
| 4 | 38766285 | G | A | G | A | 0.96 | 0.58 | 0.984 | 0.2 | 0.919 | 0.712 | 0.773 | 0.405 | 0.941 | 0.66 |
| 4 | 38767749 | G | A | G | A | 0.96 | 0.58 | 0.984 | 0.2 | 0.919 | 0.712 | 0.773 | 0.405 | 0.941 | 0.66 |
| 4 | 38769018 | A | G | A | G | 0.96 | 0.58 | 0.984 | 0.2 | 0.919 | 0.712 | 0.773 | 0.405 | 0.941 | 0.66 |
| 4 | 38777471 | C | T | C | C | 0.96 | 0.58 | 0.984 | 0.2 | 0.919 | 0.712 | 0.773 | 0.405 | 0.941 | 0.66 |
| 4 | 38778903 | G | A | G | G | 0.96 | 0.58 | 0.984 | 0.2 | 0.919 | 0.712 | 0.773 | 0.405 | 0.941 | 0.66 |
| 4 | 38779512 | C | T | C | T | 0.96 | 0.58 | 0.984 | 0.2 | 0.919 | 0.712 | 0.773 | 0.405 | 0.941 | 0.66 |
| 4 | 38780471 | T | C | T | C | 0.96 | 0.58 | 0.984 | 0.2 | 0.919 | 0.712 | 0.773 | 0.405 | 0.941 | 0.66 |
| 4 | 38783848 | T | C | C | C | 0.977 | 0.366 | 0.984 | 0.178 | 0.937 | 0.657 | 0.936 | 0.238 | 0.966 | 0.855 |
| 4 | 38789675 | G | A | A | A | 0.977 | 0.58 | 0.989 | 0.178 | 0.937 | 0.71 | 0.936 | 0.238 | 0.978 | 0.855 |
| 4 | 38796255 | G | T | T | G | 0.962 | 0.366 | 0.978 | 0.14 | 0.939 | 0.618 | 0.898 | 0.293 | 0.952 | 0.86 |
| 4 | 38802913 | A | G | G | G | 0.989 | 0.366 | 0.991 | 0.321 | 0.954 | 0.686 | 0.97 | 0.172 | 0.981 | 0.875 |
| 4 | 38803063 | A | G | G | G | 0.989 | 0.366 | 0.991 | 0.321 | 0.954 | 0.686 | 0.97 | 0.172 | 0.981 | 0.875 |
| 4 | 38803646 | T | C | C | C | 0.982 | 0.417 | 0.991 | 0.188 | 0.954 | 0.686 | 0.949 | 0.172 | 0.981 | 0.875 |
| 4 | 38803784 | C | T | T | T | 0.989 | 0.366 | 0.991 | 0.321 | 0.954 | 0.686 | 0.97 | 0.172 | 0.981 | 0.875 |
| 4 | 38804405 | C | T | T | T | 0.989 | 0.366 | 0.991 | 0.321 | 0.954 | 0.686 | 0.97 | 0.172 | 0.981 | 0.875 |
| 4 | 38805489 | C | T | T | T | 0.989 | 0.366 | 0.991 | 0.321 | 0.954 | 0.686 | 0.97 | 0.172 | 0.981 | 0.875 |
| 4 | 38805983 | T | C | C | C | 0.989 | 0.366 | 0.991 | 0.321 | 0.954 | 0.686 | 0.97 | 0.172 | 0.981 | 0.875 |
| 4 | 38806019 | A | G | G | G | 0.989 | 0.366 | 0.991 | 0.321 | 0.954 | 0.686 | 0.97 | 0.172 | 0.981 | 0.875 |
| 4 | 38806096 | T | C | C | C | 0.989 | 0.366 | 0.991 | 0.321 | 0.954 | 0.686 | 0.97 | 0.172 | 0.981 | 0.875 |
| 4 | 38806200 | G | T | T | T | 0.989 | 0.366 | 0.991 | 0.321 | 0.954 | 0.686 | 0.97 | 0.172 | 0.981 | 0.875 |
| 4 | 38806827 | G | A | A | A | 0.989 | 0.366 | 0.991 | 0.321 | 0.954 | 0.686 | 0.97 | 0.172 | 0.981 | 0.875 |
| 4 | 38806851 | A | G | G | G | 0.989 | 0.366 | 0.991 | 0.321 | 0.954 | 0.686 | 0.97 | 0.172 | 0.981 | 0.875 |
| 4 | 38807328 | A | G | G | G | 0.989 | 0.366 | 0.991 | 0.321 | 0.954 | 0.686 | 0.97 | 0.172 | 0.981 | 0.875 |
| 4 | 38808045 | A | T | T | T | 0.983 | 0.366 | 0.986 | 0.178 | 0.955 | 0.657 | 0.97 | 0.028 | 0.981 | 0.927 |
| 4 | 38808585 | G | C | C | C | 0.983 | 0.366 | 0.986 | 0.178 | 0.955 | 0.657 | 0.97 | 0.028 | 0.981 | 0.927 |
| 4 | 38808613 | G | A | A | A | 0.983 | 0.366 | 0.986 | 0.178 | 0.955 | 0.657 | 0.97 | 0.028 | 0.981 | 0.927 |
| 4 | 38809924 | G | A | A | A | 0.983 | 0.669 | 0.995 | 0.178 | 0.955 | 0.812 | 0.97 | 0.028 | 0.994 | 0.927 |
| 4 | 38810444 | C | T | T | T | 0.983 | 0.58 | 0.991 | 0.178 | 0.955 | 0.71 | 0.97 | 0.028 | 0.988 | 0.927 |
| 4 | 38810738 | C | G | G | G | 0.983 | 0.366 | 0.986 | 0.178 | 0.955 | 0.657 | 0.97 | 0.028 | 0.981 | 0.927 |
| 4 | 38810769 | C | T | T | T | 0.983 | 0.366 | 0.986 | 0.178 | 0.955 | 0.657 | 0.97 | 0.028 | 0.981 | 0.927 |
| 4 | 38810975 | C | T | T | T | 0.983 | 0.366 | 0.986 | 0.178 | 0.955 | 0.657 | 0.97 | 0.028 | 0.981 | 0.927 |
| 4 | 38811529 | T | A | A | A | 0.983 | 0.366 | 0.986 | 0.178 | 0.955 | 0.657 | 0.97 | 0.028 | 0.981 | 0.927 |
| 4 | 38811668 | G | A | A | A | 0.983 | 0.366 | 0.986 | 0.178 | 0.955 | 0.657 | 0.97 | 0.028 | 0.981 | 0.927 |
| 4 | 38811845 | G | C | C | C | 0.983 | 0.366 | 0.986 | 0.178 | 0.955 | 0.657 | 0.97 | 0.028 | 0.981 | 0.927 |
| 4 | 38812116 | T | A | A | A | 0.983 | 0.366 | 0.986 | 0.178 | 0.955 | 0.657 | 0.97 | 0.028 | 0.981 | 0.927 |
| 4 | 38812537 | A | G | G | G | 0.983 | 0.366 | 0.986 | 0.178 | 0.955 | 0.657 | 0.97 | 0.028 | 0.981 | 0.927 |
| 4 | 38814700 | A | G | G | G | 0.993 | 0.366 | 0.993 | 0.178 | 0.974 | 0.657 | 0.988 | 0.022 | 0.991 | 0.962 |
| 4 | 38827445 | A | A | G | A | - | - | - | - | - | - | - | - | - | - |
| 4 | 38828383 | A | A | G | A | - | - | - | - | - | - | - | - | - | - |
| 4 | 38833173 | A | T | T | A | 0.995 | 0.66 | 0.998 | 0.242 | 0.983 | 0.808 | 0.988 | 0.17 | 0.996 | 0.961 |
| 4 | 38835702 | T | C | C | T | 0.995 | 0.66 | 0.998 | 0.242 | 0.983 | 0.808 | 0.988 | 0.17 | 0.996 | 0.961 |
| 4 | 38836071 | G | A | A | G | 0.995 | 0.66 | 0.998 | 0.242 | 0.983 | 0.808 | 0.988 | 0.17 | 0.996 | 0.961 |
| 4 | 38843617 | A | A | T | A | - | - | - | - | - | - | - | - | - | - |
| 4 | 38843965 | T | T | C | T | - | - | - | - | - | - | - | - | - | - |
| 4 | 38846188 | T | T | G | G | - | - | - | - | - | - | - | - | - | - |
| 4 | 38851921 | T | T | G | T | - | - | - | - | - | - | - | - | - | - |
| 4 | 38853671 | G | G | A | A | - | - | - | - | - | - | - | - | - | - |
| 4 | 38856365 | T | T | C | T | - | - | - | - | - | - | - | - | - | - |
| 4 | 38857896 | C | C | A | A | - | - | - | - | - | - | - | - | - | - |
| 4 | 38871385 | T | C | C | T | 0.998 | 0.66 | 0.999 | 0.336 | 0.986 | 0.872 | 0.988 | 0.384 | 0.996 | 0.938 |
| 4 | 38876997 | G | G | A | G | - | - | - | - | - | - | - | - | - | - |
| 4 | 38879003 | A | A | C | A | - | - | - | - | - | - | - | - | - | - |
| 4 | 38881761 | G | A | A | G | 0.997 | 0.66 | 0.998 | 0.14 | 0.997 | 0.628 | 0.949 | 0.486 | 0.988 | 0.965 |
| 4 | 38881766 | G | A | A | G | 0.997 | 0.66 | 0.998 | 0.14 | 0.997 | 0.628 | 0.949 | 0.486 | 0.988 | 0.965 |
| 4 | 38881773 | A | G | G | A | 0.998 | 0.66 | 0.999 | 0.098 | 0.997 | 0.667 | 0.949 | 0.735 | 0.988 | 0.946 |
| 4 | 38886420 | G | A | A | G | 0.998 | 0.66 | 0.999 | 0.242 | 0.992 | 0.808 | 0.988 | 0.395 | 0.996 | 0.962 |
| 4 | 38892649 | T | C | C | C | 0.997 | 0.66 | 0.998 | 0.242 | 0.987 | 0.808 | 0.993 | 0.214 | 0.997 | 0.973 |
| 4 | 38894579 | C | C | T | C | - | - | - | - | - | - | - | - | - | - |
| 4 | 38905731 | T | G | G | T | 0.997 | 0.66 | 0.998 | 0.242 | 0.987 | 0.808 | 0.993 | 0.214 | 0.997 | 0.973 |

**Supplementary Table 6c:** Quantiles (1-P-values) for F_st_ values for pair-wise comparisons between Asian populations.

| chr | position | core haplotype |  |  |  | population comparison |  |  |  |  |  |  |  |  |
| --- | --- | --- | --- | --- | --- | --- | --- | --- | --- | --- | --- | --- | --- | --- |
|  |  | V | III | IV | VII | JPT<br>CHB | CHS<br>CHB | CHS<br>JPT | CDX<br>CHB | CDX<br>JPT | CDX<br>CHS | KHV<br>CHB | KHV<br>JPT | KHV<br>CHS |
| 4 | 38760338 | G | T | G | G | 0.843 | 0.942 | 0.996 | 1 | 1 | 0.997 | 1 | 1 | 0.985 |
| 4 | 38765046 | A | G | A | G | 0.926 | 0.886 | 0.997 | 1 | 1 | 0.999 | 0.999 | 1 | 0.969 |
| 4 | 38766285 | G | A | G | A | 0.926 | 0.886 | 0.997 | 1 | 1 | 0.999 | 0.999 | 1 | 0.969 |
| 4 | 38767749 | G | A | G | A | 0.926 | 0.886 | 0.997 | 1 | 1 | 0.999 | 0.999 | 1 | 0.969 |
| 4 | 38769018 | A | G | A | G | 0.926 | 0.886 | 0.997 | 1 | 1 | 0.999 | 0.999 | 1 | 0.969 |
| 4 | 38777471 | C | T | C | C | 0.879 | 0.806 | 0.99 | 1 | 1 | 0.997 | 0.996 | 0.999 | 0.93 |
| 4 | 38778903 | G | A | G | G | 0.879 | 0.806 | 0.99 | 1 | 1 | 0.997 | 0.996 | 0.999 | 0.93 |
| 4 | 38779512 | C | T | C | T | 0.879 | 0.806 | 0.99 | 1 | 1 | 0.997 | 0.996 | 0.999 | 0.93 |
| 4 | 38780471 | T | C | T | C | 0.879 | 0.806 | 0.99 | 1 | 1 | 0.997 | 0.996 | 0.999 | 0.93 |
| 4 | 38783848 | T | C | C | C | 0.642 | 0.74 | 0.924 | 0.997 | 0.999 | 0.979 | 0.993 | 0.996 | 0.925 |
| 4 | 38789675 | G | A | A | A | 0.642 | 0.74 | 0.924 | 0.997 | 0.999 | 0.979 | 0.993 | 0.996 | 0.925 |
| 4 | 38796255 | G | T | T | G | 0.358 | 0.683 | 0.822 | 0.996 | 0.996 | 0.979 | 0.993 | 0.993 | 0.943 |
| 4 | 38802913 | A | G | G | G | 0.679 | 0.683 | 0.925 | 0.999 | 1 | 0.995 | 0.995 | 0.998 | 0.96 |
| 4 | 38803063 | A | G | G | G | 0.609 | 0.742 | 0.91 | 1 | 1 | 0.998 | 0.998 | 0.998 | 0.981 |
| 4 | 38803646 | T | C | C | C | 0.679 | 0.683 | 0.925 | 0.999 | 1 | 0.995 | 0.995 | 0.998 | 0.96 |
| 4 | 38803784 | C | T | T | T | 0.679 | 0.683 | 0.925 | 0.999 | 1 | 0.995 | 0.995 | 0.998 | 0.96 |
| 4 | 38804405 | C | T | T | T | 0.679 | 0.683 | 0.925 | 0.999 | 1 | 0.995 | 0.995 | 0.998 | 0.96 |
| 4 | 38805489 | C | T | T | T | 0.679 | 0.683 | 0.925 | 0.999 | 1 | 0.995 | 0.995 | 0.998 | 0.96 |
| 4 | 38805983 | T | C | C | C | 0.679 | 0.683 | 0.925 | 0.999 | 1 | 0.995 | 0.991 | 0.997 | 0.938 |
| 4 | 38806019 | A | G | G | G | 0.679 | 0.683 | 0.925 | 0.999 | 1 | 0.995 | 0.991 | 0.997 | 0.938 |
| 4 | 38806096 | T | C | C | C | 0.679 | 0.683 | 0.925 | 0.999 | 1 | 0.995 | 0.995 | 0.998 | 0.96 |
| 4 | 38806200 | G | T | T | T | 0.679 | 0.683 | 0.925 | 0.999 | 1 | 0.995 | 0.995 | 0.998 | 0.96 |
| 4 | 38806827 | G | A | A | A | 0.78 | 0.48 | 0.925 | 0.998 | 1 | 0.995 | 0.992 | 0.998 | 0.96 |
| 4 | 38806851 | A | G | G | G | 0.78 | 0.48 | 0.925 | 0.998 | 1 | 0.995 | 0.992 | 0.998 | 0.96 |
| 4 | 38807328 | A | G | G | G | 0.78 | 0.683 | 0.944 | 0.999 | 1 | 0.995 | 0.995 | 0.998 | 0.96 |
| 4 | 38808045 | A | T | T | T | 0.78 | 0.683 | 0.944 | 0.999 | 1 | 0.995 | 0.995 | 0.998 | 0.96 |
| 4 | 38808585 | G | C | C | C | 0.78 | 0.683 | 0.944 | 0.999 | 1 | 0.995 | 0.995 | 0.998 | 0.96 |
| 4 | 38808613 | G | A | A | A | 0.679 | 0.683 | 0.925 | 0.999 | 1 | 0.995 | 0.995 | 0.998 | 0.96 |
| 4 | 38809924 | G | A | A | A | 0.679 | 0.683 | 0.925 | 0.999 | 1 | 0.995 | 0.995 | 0.998 | 0.96 |
| 4 | 38810444 | C | T | T | T | 0.78 | 0.683 | 0.944 | 0.999 | 1 | 0.995 | 0.995 | 0.998 | 0.96 |
| 4 | 38810738 | C | G | G | G | 0.78 | 0.683 | 0.944 | 0.999 | 1 | 0.995 | 0.995 | 0.998 | 0.96 |
| 4 | 38810769 | C | T | T | T | 0.78 | 0.683 | 0.944 | 0.999 | 1 | 0.995 | 0.995 | 0.998 | 0.96 |
| 4 | 38810975 | C | T | T | T | 0.78 | 0.683 | 0.944 | 0.999 | 1 | 0.995 | 0.995 | 0.998 | 0.96 |
| 4 | 38811529 | T | A | A | A | 0.78 | 0.683 | 0.944 | 0.999 | 1 | 0.995 | 0.995 | 0.998 | 0.96 |
| 4 | 38811668 | G | A | A | A | 0.78 | 0.683 | 0.944 | 0.999 | 1 | 0.995 | 0.995 | 0.998 | 0.96 |
| 4 | 38811845 | G | C | C | C | 0.78 | 0.683 | 0.944 | 0.999 | 1 | 0.995 | 0.995 | 0.998 | 0.96 |
| 4 | 38812116 | T | A | A | A | 0.78 | 0.683 | 0.944 | 0.999 | 1 | 0.995 | 0.995 | 0.998 | 0.96 |
| 4 | 38812537 | A | G | G | G | 0.78 | 0.683 | 0.944 | 0.999 | 1 | 0.995 | 0.995 | 0.998 | 0.96 |
| 4 | 38814700 | A | G | G | G | 0.679 | 0.775 | 0.956 | 0.999 | 1 | 0.992 | 0.996 | 0.998 | 0.941 |
| 4 | 38827445 | A | A | G | A | 0.69 | 0.384 | 0.177 | 0.679 | 0.952 | 0.963 | 0.019 | 0.632 | 0.291 |
| 4 | 38828383 | A | A | G | A | 0.69 | 0.384 | 0.177 | 0.679 | 0.952 | 0.963 | 0.019 | 0.632 | 0.291 |
| 4 | 38833173 | A | T | T | A | 0.676 | 0.874 | 0.968 | 0.993 | 0.998 | 0.904 | 0.999 | 0.999 | 0.968 |
| 4 | 38835702 | T | C | C | T | 0.719 | 0.874 | 0.979 | 0.993 | 0.999 | 0.904 | 0.999 | 0.999 | 0.968 |
| 4 | 38836071 | G | A | A | G | 0.719 | 0.874 | 0.979 | 0.993 | 0.999 | 0.904 | 0.999 | 0.999 | 0.968 |
| 4 | 38843617 | A | A | T | A | 0.69 | 0.384 | 0.177 | 0.679 | 0.952 | 0.963 | 0.019 | 0.632 | 0.291 |
| 4 | 38843965 | T | T | C | T | 0.69 | 0.384 | 0.177 | 0.679 | 0.952 | 0.963 | 0.019 | 0.632 | 0.291 |
| 4 | 38846188 | T | T | G | G | 0.69 | 0.384 | 0.177 | 0.679 | 0.952 | 0.963 | 0.019 | 0.632 | 0.291 |
| 4 | 38851921 | T | T | G | T | 0.69 | 0.384 | 0.177 | 0.679 | 0.952 | 0.963 | 0.019 | 0.632 | 0.291 |
| 4 | 38853671 | G | G | A | A | 0.69 | 0.384 | 0.177 | 0.679 | 0.952 | 0.963 | 0.019 | 0.632 | 0.291 |
| 4 | 38856365 | T | T | C | T | 0.69 | 0.384 | 0.177 | 0.311 | 0.929 | 0.939 | 0.019 | 0.632 | 0.291 |
| 4 | 38857896 | C | C | A | A | 0.69 | 0.384 | 0.177 | 0.311 | 0.929 | 0.939 | 0.019 | 0.632 | 0.291 |
| 4 | 38871385 | T | C | C | T | 0.643 | 0.893 | 0.974 | 0.991 | 0.996 | 0.832 | 0.996 | 0.998 | 0.894 |
| 4 | 38876997 | G | G | A | G | 0.69 | 0.384 | 0.177 | 0.311 | 0.929 | 0.939 | 0.019 | 0.632 | 0.291 |
| 4 | 38879003 | A | A | C | A | 0.69 | 0.384 | 0.177 | 0.311 | 0.929 | 0.939 | 0.019 | 0.632 | 0.291 |
| 4 | 38881761 | G | A | A | G | 0.718 | 0.902 | 0.989 | 0.985 | 0.997 | 0.707 | 0.987 | 0.997 | 0.702 |
| 4 | 38881766 | G | A | A | G | 0.718 | 0.902 | 0.989 | 0.985 | 0.997 | 0.707 | 0.987 | 0.997 | 0.702 |
| 4 | 38881773 | A | G | G | A | 0.813 | 0.885 | 0.989 | 0.987 | 0.998 | 0.756 | 0.983 | 0.997 | 0.702 |
| 4 | 38886420 | G | A | A | G | 0.139 | 0.938 | 0.91 | 0.994 | 0.99 | 0.836 | 0.996 | 0.991 | 0.868 |
| 4 | 38892649 | T | C | C | C | 0.139 | 0.938 | 0.91 | 0.994 | 0.99 | 0.836 | 0.996 | 0.991 | 0.868 |
| 4 | 38894579 | C | C | T | C | 0.69 | 0.384 | 0.177 | 0.311 | 0.929 | 0.939 | 0.019 | 0.632 | 0.291 |
| 4 | 38905731 | T | G | G | T | 0.139 | 0.954 | 0.925 | 0.994 | 0.99 | 0.752 | 0.996 | 0.991 | 0.84 |
|  |  | V | III | IV | VII | KHV<br>CDX | GIH<br>CHB | GIH<br>JPT | GIH<br>CHS | GIH<br>CDX | GIH<br>KHV | PJL<br>CHB | PJL<br>JPT | PJL<br>CHS |
| 4 | 38760338 | G | T | G | G | 0.395 | 0.505 | 0.883 | 0.045 | 0.906 | 0.843 | 0.933 | 0.988 | 0.616 |
| 4 | 38765046 | A | G | A | G | 0.757 | 0.49 | 0.918 | 0.046 | 0.916 | 0.766 | 0.927 | 0.991 | 0.643 |
| 4 | 38766285 | G | A | G | A | 0.757 | 0.49 | 0.918 | 0.046 | 0.916 | 0.766 | 0.927 | 0.991 | 0.643 |
| 4 | 38767749 | G | A | G | A | 0.757 | 0.511 | 0.929 | 0.044 | 0.903 | 0.723 | 0.941 | 0.992 | 0.718 |
| 4 | 38769018 | A | G | A | G | 0.757 | 0.511 | 0.929 | 0.044 | 0.903 | 0.723 | 0.941 | 0.992 | 0.718 |
| 4 | 38777471 | C | T | C | C | 0.763 | 0.629 | 0.941 | 0.108 | 0.75 | 0.391 | 0.962 | 0.993 | 0.851 |
| 4 | 38778903 | G | A | G | G | 0.763 | 0.629 | 0.941 | 0.108 | 0.75 | 0.391 | 0.962 | 0.993 | 0.851 |
| 4 | 38779512 | C | T | C | T | 0.763 | 0.629 | 0.941 | 0.108 | 0.75 | 0.391 | 0.962 | 0.993 | 0.851 |
| 4 | 38780471 | T | C | T | C | 0.763 | 0.629 | 0.941 | 0.108 | 0.75 | 0.391 | 0.962 | 0.993 | 0.851 |
| 4 | 38783848 | T | C | C | C | 0.368 | 0.9 | 0.967 | 0.68 | 0.047 | 0.054 | 0.985 | 0.994 | 0.952 |
| 4 | 38789675 | G | A | A | A | 0.368 | 0.9 | 0.967 | 0.68 | 0.047 | 0.054 | 0.985 | 0.994 | 0.952 |
| 4 | 38796255 | G | T | T | G | 0.284 | 0.905 | 0.952 | 0.738 | 0.043 | 0.084 | 0.982 | 0.991 | 0.952 |
| 4 | 38802913 | A | G | G | G | 0.423 | 0.925 | 0.979 | 0.786 | 0.047 | 0.086 | 0.988 | 0.997 | 0.969 |
| 4 | 38803063 | A | G | G | G | 0.423 | 0.952 | 0.982 | 0.838 | 0.047 | 0.086 | 0.992 | 0.997 | 0.976 |
| 4 | 38803646 | T | C | C | C | 0.423 | 0.925 | 0.979 | 0.786 | 0.047 | 0.086 | 0.988 | 0.997 | 0.969 |
| 4 | 38803784 | C | T | T | T | 0.423 | 0.925 | 0.979 | 0.786 | 0.047 | 0.086 | 0.988 | 0.997 | 0.969 |
| 4 | 38804405 | C | T | T | T | 0.423 | 0.925 | 0.979 | 0.786 | 0.047 | 0.086 | 0.988 | 0.997 | 0.969 |
| 4 | 38805489 | C | T | T | T | 0.423 | 0.925 | 0.979 | 0.786 | 0.047 | 0.086 | 0.988 | 0.997 | 0.969 |
| 4 | 38805983 | T | C | C | C | 0.469 | 0.925 | 0.979 | 0.786 | 0.047 | 0.116 | 0.986 | 0.996 | 0.962 |
| 4 | 38806019 | A | G | G | G | 0.469 | 0.925 | 0.979 | 0.786 | 0.047 | 0.116 | 0.986 | 0.996 | 0.962 |
| 4 | 38806096 | T | C | C | C | 0.423 | 0.925 | 0.979 | 0.786 | 0.047 | 0.086 | 0.988 | 0.997 | 0.969 |
| 4 | 38806200 | G | T | T | T | 0.423 | 0.925 | 0.979 | 0.786 | 0.047 | 0.086 | 0.988 | 0.997 | 0.969 |
| 4 | 38806827 | G | A | A | A | 0.423 | 0.916 | 0.979 | 0.786 | 0.047 | 0.086 | 0.987 | 0.997 | 0.969 |
| 4 | 38806851 | A | G | G | G | 0.423 | 0.916 | 0.979 | 0.786 | 0.047 | 0.086 | 0.987 | 0.997 | 0.969 |
| 4 | 38807328 | A | G | G | G | 0.423 | 0.925 | 0.982 | 0.786 | 0.047 | 0.086 | 0.988 | 0.997 | 0.969 |
| 4 | 38808045 | A | T | T | T | 0.423 | 0.925 | 0.982 | 0.786 | 0.047 | 0.086 | 0.988 | 0.997 | 0.969 |
| 4 | 38808585 | G | C | C | C | 0.423 | 0.925 | 0.982 | 0.786 | 0.047 | 0.086 | 0.988 | 0.997 | 0.969 |
| 4 | 38808613 | G | A | A | A | 0.423 | 0.925 | 0.979 | 0.786 | 0.047 | 0.086 | 0.988 | 0.997 | 0.969 |
| 4 | 38809924 | G | A | A | A | 0.423 | 0.925 | 0.979 | 0.786 | 0.047 | 0.086 | 0.988 | 0.997 | 0.969 |
| 4 | 38810444 | C | T | T | T | 0.423 | 0.925 | 0.982 | 0.786 | 0.047 | 0.086 | 0.988 | 0.997 | 0.969 |
| 4 | 38810738 | C | G | G | G | 0.423 | 0.925 | 0.982 | 0.786 | 0.047 | 0.086 | 0.988 | 0.997 | 0.969 |
| 4 | 38810769 | C | T | T | T | 0.423 | 0.925 | 0.982 | 0.786 | 0.047 | 0.086 | 0.988 | 0.997 | 0.969 |
| 4 | 38810975 | C | T | T | T | 0.423 | 0.925 | 0.982 | 0.786 | 0.047 | 0.086 | 0.988 | 0.997 | 0.969 |
| 4 | 38811529 | T | A | A | A | 0.423 | 0.925 | 0.982 | 0.786 | 0.047 | 0.086 | 0.988 | 0.997 | 0.969 |
| 4 | 38811668 | G | A | A | A | 0.423 | 0.925 | 0.982 | 0.786 | 0.047 | 0.086 | 0.988 | 0.997 | 0.969 |
| 4 | 38811845 | G | C | C | C | 0.423 | 0.925 | 0.982 | 0.786 | 0.047 | 0.086 | 0.988 | 0.997 | 0.969 |
| 4 | 38812116 | T | A | A | A | 0.423 | 0.925 | 0.982 | 0.786 | 0.047 | 0.086 | 0.988 | 0.997 | 0.969 |
| 4 | 38812537 | A | G | G | G | 0.423 | 0.925 | 0.982 | 0.786 | 0.047 | 0.086 | 0.988 | 0.997 | 0.969 |
| 4 | 38814700 | A | G | G | G | 0.423 | 0.868 | 0.965 | 0.6 | 0.327 | 0.053 | 0.982 | 0.995 | 0.936 |
| 4 | 38827445 | A | A | G | A | 0.918 | 0.925 | 0.746 | 0.84 | 0.979 | 0.924 | 0.865 | 0.463 | 0.508 |
| 4 | 38828383 | A | A | G | A | 0.918 | 0.925 | 0.746 | 0.84 | 0.979 | 0.924 | 0.865 | 0.463 | 0.508 |
| 4 | 38833173 | A | T | T | A | 0.176 | 0.678 | 0.901 | 0.103 | 0.351 | 0.532 | 0.946 | 0.984 | 0.758 |
| 4 | 38835702 | T | C | C | T | 0.176 | 0.678 | 0.916 | 0.103 | 0.351 | 0.532 | 0.946 | 0.986 | 0.758 |
| 4 | 38836071 | G | A | A | G | 0.176 | 0.678 | 0.916 | 0.103 | 0.351 | 0.532 | 0.946 | 0.986 | 0.758 |
| 4 | 38843617 | A | A | T | A | 0.918 | 0.925 | 0.746 | 0.84 | 0.979 | 0.924 | 0.865 | 0.463 | 0.508 |
| 4 | 38843965 | T | T | C | T | 0.918 | 0.925 | 0.746 | 0.84 | 0.979 | 0.924 | 0.865 | 0.463 | 0.508 |
| 4 | 38846188 | T | T | G | G | 0.918 | 0.925 | 0.746 | 0.84 | 0.979 | 0.924 | 0.865 | 0.463 | 0.508 |
| 4 | 38851921 | T | T | G | T | 0.918 | 0.925 | 0.746 | 0.84 | 0.979 | 0.924 | 0.865 | 0.463 | 0.508 |
| 4 | 38853671 | G | G | A | A | 0.918 | 0.925 | 0.746 | 0.84 | 0.979 | 0.924 | 0.865 | 0.463 | 0.508 |
| 4 | 38856365 | T | T | C | T | 0.793 | 0.925 | 0.746 | 0.84 | 0.972 | 0.924 | 0.865 | 0.463 | 0.508 |
| 4 | 38857896 | C | C | A | A | 0.793 | 0.925 | 0.746 | 0.84 | 0.972 | 0.924 | 0.865 | 0.463 | 0.508 |
| 4 | 38871385 | T | C | C | T | 0.139 | 0.678 | 0.889 | 0.078 | 0.136 | 0.388 | 0.924 | 0.975 | 0.622 |
| 4 | 38876997 | G | G | A | G | 0.793 | 0.925 | 0.746 | 0.84 | 0.972 | 0.924 | 0.683 | 0.062 | 0.111 |
| 4 | 38879003 | A | A | C | A | 0.793 | 0.925 | 0.746 | 0.84 | 0.972 | 0.924 | 0.683 | 0.062 | 0.111 |
| 4 | 38881761 | G | A | A | G | 0.085 | 0.629 | 0.914 | 0.045 | 0.12 | 0.123 | 0.868 | 0.968 | 0.368 |
| 4 | 38881766 | G | A | A | G | 0.085 | 0.629 | 0.914 | 0.045 | 0.12 | 0.123 | 0.868 | 0.968 | 0.368 |
| 4 | 38881773 | A | G | G | A | 0.165 | 0.607 | 0.914 | 0.045 | 0.327 | 0.123 | 0.81 | 0.963 | 0.35 |
| 4 | 38886420 | G | A | A | G | 0.153 | 0.738 | 0.782 | 0.078 | 0.342 | 0.372 | 0.924 | 0.942 | 0.543 |
| 4 | 38892649 | T | C | C | C | 0.153 | 0.738 | 0.782 | 0.078 | 0.342 | 0.372 | 0.924 | 0.942 | 0.543 |
| 4 | 38894579 | C | C | T | C | 0.793 | 0.925 | 0.746 | 0.84 | 0.972 | 0.924 | 0.683 | 0.062 | 0.111 |
| 4 | 38905731 | T | G | G | T | 0.153 | 0.738 | 0.782 | 0.056 | 0.342 | 0.372 | 0.924 | 0.942 | 0.52 |
|  |  | V | III | IV | VII | PJL<br>CDX | PJL<br>KHV | PJL<br>GIH | BEB<br>CHB | BEB<br>JPT | BEB<br>CHS | BEB<br>CDX | BEB<br>KHV | BEB<br>GIH |
| 4 | 38760338 | G | T | G | G | 0.378 | 0.121 | 0.96 | 0.448 | 0.903 | 0.097 | 0.95 | 0.901 | 0.026 |
| 4 | 38765046 | A | G | A | G | 0.398 | 0.054 | 0.964 | 0.673 | 0.964 | 0.068 | 0.924 | 0.765 | 0.259 |
| 4 | 38766285 | G | A | G | A | 0.398 | 0.054 | 0.964 | 0.673 | 0.964 | 0.068 | 0.924 | 0.765 | 0.259 |
| 4 | 38767749 | G | A | G | A | 0.372 | 0.055 | 0.971 | 0.773 | 0.979 | 0.149 | 0.857 | 0.6 | 0.38 |
| 4 | 38769018 | A | G | A | G | 0.372 | 0.055 | 0.971 | 0.773 | 0.979 | 0.149 | 0.857 | 0.6 | 0.38 |
| 4 | 38777471 | C | T | C | C | 0.048 | 0.381 | 0.971 | 0.858 | 0.982 | 0.443 | 0.671 | 0.154 | 0.38 |
| 4 | 38778903 | G | A | G | G | 0.048 | 0.381 | 0.971 | 0.858 | 0.982 | 0.443 | 0.671 | 0.154 | 0.38 |
| 4 | 38779512 | C | T | C | T | 0.048 | 0.381 | 0.971 | 0.858 | 0.982 | 0.443 | 0.671 | 0.154 | 0.38 |
| 4 | 38780471 | T | C | T | C | 0.048 | 0.381 | 0.971 | 0.858 | 0.982 | 0.443 | 0.671 | 0.154 | 0.38 |
| 4 | 38783848 | T | C | C | C | 0.522 | 0.728 | 0.929 | 0.962 | 0.988 | 0.855 | 0.053 | 0.19 | 0.33 |
| 4 | 38789675 | G | A | A | A | 0.522 | 0.728 | 0.929 | 0.962 | 0.988 | 0.855 | 0.053 | 0.19 | 0.33 |
| 4 | 38796255 | G | T | T | G | 0.522 | 0.719 | 0.884 | 0.957 | 0.98 | 0.855 | 0.053 | 0.172 | 0.261 |
| 4 | 38802913 | A | G | G | G | 0.522 | 0.775 | 0.929 | 0.972 | 0.993 | 0.916 | 0.053 | 0.425 | 0.33 |
| 4 | 38803063 | A | G | G | G | 0.522 | 0.775 | 0.929 | 0.983 | 0.994 | 0.938 | 0.053 | 0.425 | 0.33 |
| 4 | 38803646 | T | C | C | C | 0.522 | 0.775 | 0.929 | 0.972 | 0.993 | 0.916 | 0.053 | 0.425 | 0.33 |
| 4 | 38803784 | C | T | T | T | 0.522 | 0.775 | 0.929 | 0.972 | 0.993 | 0.916 | 0.053 | 0.425 | 0.33 |
| 4 | 38804405 | C | T | T | T | 0.522 | 0.775 | 0.929 | 0.972 | 0.993 | 0.916 | 0.053 | 0.425 | 0.33 |
| 4 | 38805489 | C | T | T | T | 0.522 | 0.775 | 0.929 | 0.972 | 0.993 | 0.916 | 0.053 | 0.425 | 0.33 |
| 4 | 38805983 | T | C | C | C | 0.378 | 0.744 | 0.868 | 0.972 | 0.993 | 0.916 | 0.053 | 0.444 | 0.33 |
| 4 | 38806019 | A | G | G | G | 0.378 | 0.744 | 0.868 | 0.972 | 0.993 | 0.916 | 0.053 | 0.444 | 0.33 |
| 4 | 38806096 | T | C | C | C | 0.522 | 0.775 | 0.929 | 0.972 | 0.993 | 0.916 | 0.053 | 0.425 | 0.33 |
| 4 | 38806200 | G | T | T | T | 0.522 | 0.775 | 0.929 | 0.972 | 0.993 | 0.916 | 0.053 | 0.425 | 0.33 |
| 4 | 38806827 | G | A | A | A | 0.522 | 0.775 | 0.929 | 0.967 | 0.993 | 0.916 | 0.053 | 0.425 | 0.33 |
| 4 | 38806851 | A | G | G | G | 0.522 | 0.775 | 0.929 | 0.967 | 0.993 | 0.916 | 0.053 | 0.425 | 0.33 |
| 4 | 38807328 | A | G | G | G | 0.522 | 0.775 | 0.929 | 0.972 | 0.994 | 0.916 | 0.053 | 0.425 | 0.33 |
| 4 | 38808045 | A | T | T | T | 0.522 | 0.775 | 0.929 | 0.972 | 0.994 | 0.916 | 0.053 | 0.425 | 0.33 |
| 4 | 38808585 | G | C | C | C | 0.522 | 0.775 | 0.929 | 0.972 | 0.994 | 0.916 | 0.053 | 0.425 | 0.33 |
| 4 | 38808613 | G | A | A | A | 0.522 | 0.775 | 0.929 | 0.972 | 0.993 | 0.916 | 0.053 | 0.425 | 0.33 |
| 4 | 38809924 | G | A | A | A | 0.522 | 0.775 | 0.929 | 0.972 | 0.993 | 0.916 | 0.053 | 0.425 | 0.33 |
| 4 | 38810444 | C | T | T | T | 0.522 | 0.775 | 0.929 | 0.972 | 0.994 | 0.916 | 0.053 | 0.425 | 0.33 |
| 4 | 38810738 | C | G | G | G | 0.522 | 0.775 | 0.929 | 0.972 | 0.994 | 0.916 | 0.053 | 0.425 | 0.33 |
| 4 | 38810769 | C | T | T | T | 0.522 | 0.775 | 0.929 | 0.972 | 0.994 | 0.916 | 0.053 | 0.425 | 0.33 |
| 4 | 38810975 | C | T | T | T | 0.522 | 0.775 | 0.929 | 0.972 | 0.994 | 0.916 | 0.053 | 0.425 | 0.33 |
| 4 | 38811529 | T | A | A | A | 0.522 | 0.775 | 0.929 | 0.972 | 0.994 | 0.916 | 0.053 | 0.425 | 0.33 |
| 4 | 38811668 | G | A | A | A | 0.522 | 0.775 | 0.929 | 0.972 | 0.994 | 0.916 | 0.053 | 0.425 | 0.33 |
| 4 | 38811845 | G | C | C | C | 0.522 | 0.775 | 0.929 | 0.972 | 0.994 | 0.916 | 0.053 | 0.425 | 0.33 |
| 4 | 38812116 | T | A | A | A | 0.522 | 0.775 | 0.929 | 0.972 | 0.994 | 0.916 | 0.053 | 0.425 | 0.33 |
| 4 | 38812537 | A | G | G | G | 0.522 | 0.775 | 0.929 | 0.972 | 0.994 | 0.916 | 0.053 | 0.425 | 0.33 |
| 4 | 38814700 | A | G | G | G | 0.14 | 0.579 | 0.935 | 0.971 | 0.993 | 0.876 | 0.044 | 0.19 | 0.681 |
| 4 | 38827445 | A | A | G | A | 0.968 | 0.822 | 0.668 | 0.949 | 0.79 | 0.874 | 0.989 | 0.942 | - |
| 4 | 38828383 | A | A | G | A | 0.968 | 0.822 | 0.668 | 0.949 | 0.79 | 0.874 | 0.989 | 0.942 | - |
| 4 | 38833173 | A | T | T | A | 0.059 | 0.053 | 0.884 | 0.933 | 0.983 | 0.67 | 0.009 | 0.112 | 0.709 |
| 4 | 38835702 | T | C | C | T | 0.059 | 0.053 | 0.884 | 0.933 | 0.985 | 0.67 | 0.009 | 0.112 | 0.709 |
| 4 | 38836071 | G | A | A | G | 0.059 | 0.053 | 0.884 | 0.933 | 0.985 | 0.67 | 0.009 | 0.112 | 0.709 |
| 4 | 38843617 | A | A | T | A | 0.968 | 0.822 | 0.668 | 0.949 | 0.79 | 0.874 | 0.989 | 0.942 | - |
| 4 | 38843965 | T | T | C | T | 0.968 | 0.822 | 0.668 | 0.949 | 0.79 | 0.874 | 0.989 | 0.942 | - |
| 4 | 38846188 | T | T | G | G | 0.968 | 0.822 | 0.668 | 0.949 | 0.79 | 0.874 | 0.989 | 0.942 | - |
| 4 | 38851921 | T | T | G | T | 0.968 | 0.822 | 0.668 | 0.949 | 0.79 | 0.874 | 0.989 | 0.942 | - |
| 4 | 38853671 | G | G | A | A | 0.968 | 0.822 | 0.668 | 0.949 | 0.79 | 0.874 | 0.989 | 0.942 | - |
| 4 | 38856365 | T | T | C | T | 0.958 | 0.822 | 0.668 | 0.949 | 0.79 | 0.874 | 0.985 | 0.942 | - |
| 4 | 38857896 | C | C | A | A | 0.958 | 0.822 | 0.668 | 0.949 | 0.79 | 0.874 | 0.985 | 0.942 | - |
| 4 | 38871385 | T | C | C | T | 0.028 | 0.054 | 0.737 | 0.899 | 0.973 | 0.443 | 0.034 | 0.113 | 0.409 |
| 4 | 38876997 | G | G | A | G | 0.925 | 0.712 | 0.869 | 0.949 | 0.79 | 0.874 | 0.985 | 0.942 | - |
| 4 | 38879003 | A | A | C | A | 0.925 | 0.712 | 0.869 | 0.949 | 0.79 | 0.874 | 0.985 | 0.942 | - |
| 4 | 38881761 | G | A | A | G | 0.002 | 0.053 | 0.416 | 0.874 | 0.975 | 0.397 | 0.007 | 0.069 | 0.359 |
| 4 | 38881766 | G | A | A | G | 0.002 | 0.053 | 0.416 | 0.874 | 0.975 | 0.397 | 0.007 | 0.069 | 0.359 |
| 4 | 38881773 | A | G | G | A | 0.008 | 0.05 | 0.348 | 0.819 | 0.97 | 0.168 | 0.101 | 0.083 | 0.261 |
| 4 | 38886420 | G | A | A | G | 0.009 | 0.054 | 0.712 | 0.899 | 0.923 | 0.414 | 0.075 | 0.113 | 0.36 |
| 4 | 38892649 | T | C | C | C | 0.009 | 0.054 | 0.712 | 0.899 | 0.923 | 0.414 | 0.075 | 0.113 | 0.36 |
| 4 | 38894579 | C | C | T | C | 0.925 | 0.712 | 0.869 | 0.949 | 0.79 | 0.874 | 0.985 | 0.942 | - |
| 4 | 38905731 | T | G | G | T | 0.009 | 0.054 | 0.712 | 0.899 | 0.923 | 0.395 | 0.075 | 0.113 | 0.36 |
|  |  | V | III | IV | VII | BEB<br>PJL | STU<br>CHB | STU<br>JPT | STU<br>CHS | STU<br>CDX | STU<br>KHV | STU<br>GIH | STU<br>PJL | STU<br>BEB |
| 4 | 38760338 | G | T | G | G | 0.972 | 0.63 | 0.929 | 0.042 | 0.885 | 0.794 | 0.147 | 0.919 | 0.229 |
| 4 | 38765046 | A | G | A | G | 0.929 | 0.811 | 0.98 | 0.366 | 0.76 | 0.41 | 0.712 | 0.444 | 0.451 |
| 4 | 38766285 | G | A | G | A | 0.929 | 0.811 | 0.98 | 0.366 | 0.76 | 0.41 | 0.712 | 0.444 | 0.451 |
| 4 | 38767749 | G | A | G | A | 0.869 | 0.915 | 0.989 | 0.624 | 0.543 | 0.096 | 0.918 | 0.272 | 0.724 |
| 4 | 38769018 | A | G | A | G | 0.869 | 0.915 | 0.989 | 0.624 | 0.543 | 0.096 | 0.918 | 0.272 | 0.724 |
| 4 | 38777471 | C | T | C | C | 0.869 | 0.955 | 0.992 | 0.817 | 0.084 | 0.154 | 0.95 | 0.218 | 0.774 |
| 4 | 38778903 | G | A | G | G | 0.869 | 0.955 | 0.992 | 0.817 | 0.084 | 0.154 | 0.95 | 0.218 | 0.774 |
| 4 | 38779512 | C | T | C | T | 0.869 | 0.944 | 0.991 | 0.777 | 0.116 | 0.128 | 0.918 | 0.272 | 0.724 |
| 4 | 38780471 | T | C | T | C | 0.869 | 0.944 | 0.991 | 0.777 | 0.116 | 0.128 | 0.918 | 0.272 | 0.724 |
| 4 | 38783848 | T | C | C | C | 0.773 | 0.978 | 0.993 | 0.932 | 0.37 | 0.652 | 0.837 | 0.273 | 0.648 |
| 4 | 38789675 | G | A | A | A | 0.773 | 0.974 | 0.991 | 0.916 | 0.145 | 0.565 | 0.716 | 0.37 | 0.411 |
| 4 | 38796255 | G | T | T | G | 0.773 | 0.97 | 0.986 | 0.916 | 0.145 | 0.549 | 0.688 | 0.37 | 0.411 |
| 4 | 38802913 | A | G | G | G | 0.773 | 0.983 | 0.995 | 0.956 | 0.37 | 0.675 | 0.837 | 0.273 | 0.648 |
| 4 | 38803063 | A | G | G | G | 0.773 | 0.99 | 0.995 | 0.967 | 0.37 | 0.675 | 0.837 | 0.273 | 0.648 |
| 4 | 38803646 | T | C | C | C | 0.773 | 0.983 | 0.995 | 0.956 | 0.37 | 0.675 | 0.837 | 0.273 | 0.648 |
| 4 | 38803784 | C | T | T | T | 0.773 | 0.983 | 0.995 | 0.956 | 0.37 | 0.675 | 0.837 | 0.273 | 0.648 |
| 4 | 38804405 | C | T | T | T | 0.773 | 0.983 | 0.995 | 0.956 | 0.37 | 0.675 | 0.837 | 0.273 | 0.648 |
| 4 | 38805489 | C | T | T | T | 0.773 | 0.983 | 0.995 | 0.956 | 0.37 | 0.675 | 0.837 | 0.273 | 0.648 |
| 4 | 38805983 | T | C | C | C | 0.712 | 0.983 | 0.995 | 0.956 | 0.37 | 0.725 | 0.837 | 0.166 | 0.648 |
| 4 | 38806019 | A | G | G | G | 0.712 | 0.983 | 0.995 | 0.956 | 0.37 | 0.725 | 0.837 | 0.166 | 0.648 |
| 4 | 38806096 | T | C | C | C | 0.773 | 0.983 | 0.995 | 0.956 | 0.37 | 0.675 | 0.837 | 0.273 | 0.648 |
| 4 | 38806200 | G | T | T | T | 0.773 | 0.983 | 0.995 | 0.956 | 0.37 | 0.675 | 0.837 | 0.273 | 0.648 |
| 4 | 38806827 | G | A | A | A | 0.773 | 0.981 | 0.995 | 0.956 | 0.37 | 0.675 | 0.837 | 0.273 | 0.648 |
| 4 | 38806851 | A | G | G | G | 0.773 | 0.981 | 0.995 | 0.956 | 0.37 | 0.675 | 0.837 | 0.273 | 0.648 |
| 4 | 38807328 | A | G | G | G | 0.773 | 0.983 | 0.996 | 0.956 | 0.37 | 0.675 | 0.837 | 0.273 | 0.648 |
| 4 | 38808045 | A | T | T | T | 0.773 | 0.983 | 0.996 | 0.956 | 0.37 | 0.675 | 0.837 | 0.273 | 0.648 |
| 4 | 38808585 | G | C | C | C | 0.773 | 0.983 | 0.996 | 0.956 | 0.37 | 0.675 | 0.837 | 0.273 | 0.648 |
| 4 | 38808613 | G | A | A | A | 0.773 | 0.983 | 0.995 | 0.956 | 0.37 | 0.675 | 0.837 | 0.273 | 0.648 |
| 4 | 38809924 | G | A | A | A | 0.773 | 0.983 | 0.995 | 0.956 | 0.37 | 0.675 | 0.837 | 0.273 | 0.648 |
| 4 | 38810444 | C | T | T | T | 0.773 | 0.983 | 0.996 | 0.956 | 0.37 | 0.675 | 0.837 | 0.273 | 0.648 |
| 4 | 38810738 | C | G | G | G | 0.773 | 0.983 | 0.996 | 0.956 | 0.37 | 0.675 | 0.837 | 0.273 | 0.648 |
| 4 | 38810769 | C | T | T | T | 0.773 | 0.983 | 0.996 | 0.956 | 0.37 | 0.675 | 0.837 | 0.273 | 0.648 |
| 4 | 38810975 | C | T | T | T | 0.773 | 0.983 | 0.996 | 0.956 | 0.37 | 0.675 | 0.837 | 0.273 | 0.648 |
| 4 | 38811529 | T | A | A | A | 0.773 | 0.983 | 0.996 | 0.956 | 0.37 | 0.675 | 0.837 | 0.273 | 0.648 |
| 4 | 38811668 | G | A | A | A | 0.773 | 0.983 | 0.996 | 0.956 | 0.37 | 0.675 | 0.837 | 0.273 | 0.648 |
| 4 | 38811845 | G | C | C | C | 0.773 | 0.983 | 0.996 | 0.956 | 0.37 | 0.675 | 0.837 | 0.273 | 0.648 |
| 4 | 38812116 | T | A | A | A | 0.773 | 0.983 | 0.996 | 0.956 | 0.37 | 0.675 | 0.837 | 0.273 | 0.648 |
| 4 | 38812537 | A | G | G | G | 0.773 | 0.983 | 0.996 | 0.956 | 0.37 | 0.675 | 0.837 | 0.273 | 0.648 |
| 4 | 38814700 | A | G | G | G | 0.449 | 0.976 | 0.993 | 0.912 | 0.088 | 0.438 | 0.853 | 0.267 | 0.371 |
| 4 | 38827445 | A | A | G | A | 0.697 | 0.852 | 0.457 | 0.508 | 0.971 | 0.819 | 0.675 | 0.096 | 0.691 |
| 4 | 38828383 | A | A | G | A | 0.697 | 0.852 | 0.457 | 0.508 | 0.971 | 0.819 | 0.675 | 0.096 | 0.691 |
| 4 | 38833173 | A | T | T | A | 0.275 | 0.947 | 0.986 | 0.763 | 0.087 | 0.091 | 0.881 | 0.142 | 0.37 |
| 4 | 38835702 | T | C | C | T | 0.275 | 0.947 | 0.987 | 0.763 | 0.087 | 0.091 | 0.881 | 0.142 | 0.37 |
| 4 | 38836071 | G | A | A | G | 0.275 | 0.947 | 0.987 | 0.763 | 0.087 | 0.091 | 0.881 | 0.142 | 0.37 |
| 4 | 38843617 | A | A | T | A | 0.697 | 0.852 | 0.457 | 0.508 | 0.971 | 0.819 | 0.675 | 0.096 | 0.691 |
| 4 | 38843965 | T | T | C | T | 0.697 | 0.852 | 0.457 | 0.508 | 0.971 | 0.819 | 0.675 | 0.096 | 0.691 |
| 4 | 38846188 | T | T | G | G | 0.697 | 0.687 | 0.053 | 0.101 | 0.943 | 0.708 | 0.87 | 0.289 | 0.895 |
| 4 | 38851921 | T | T | G | T | 0.697 | 0.852 | 0.457 | 0.508 | 0.971 | 0.819 | 0.675 | 0.096 | 0.691 |
| 4 | 38853671 | G | G | A | A | 0.697 | 0.687 | 0.053 | 0.101 | 0.943 | 0.708 | 0.87 | 0.289 | 0.895 |
| 4 | 38856365 | T | T | C | T | 0.697 | 0.852 | 0.457 | 0.508 | 0.958 | 0.819 | 0.675 | 0.096 | 0.691 |
| 4 | 38857896 | C | C | A | A | 0.697 | 0.687 | 0.053 | 0.101 | 0.92 | 0.708 | 0.87 | 0.289 | 0.895 |
| 4 | 38871385 | T | C | C | T | 0.275 | 0.94 | 0.981 | 0.701 | 0.087 | 0.097 | 0.853 | 0.161 | 0.411 |
| 4 | 38876997 | G | G | A | G | 0.897 | 0.852 | 0.457 | 0.508 | 0.958 | 0.819 | 0.675 | 0.289 | 0.691 |
| 4 | 38879003 | A | A | C | A | 0.897 | 0.852 | 0.457 | 0.508 | 0.958 | 0.819 | 0.675 | 0.289 | 0.691 |
| 4 | 38881761 | G | A | A | G | 0.087 | 0.894 | 0.976 | 0.52 | 0.012 | 0.096 | 0.679 | 0.154 | 0.278 |
| 4 | 38881766 | G | A | A | G | 0.087 | 0.894 | 0.976 | 0.52 | 0.012 | 0.096 | 0.679 | 0.154 | 0.278 |
| 4 | 38881773 | A | G | G | A | 0.129 | 0.875 | 0.976 | 0.52 | 0.009 | 0.096 | 0.679 | 0.218 | 0.37 |
| 4 | 38886420 | G | A | A | G | 0.275 | 0.94 | 0.956 | 0.642 | 0.07 | 0.097 | 0.825 | 0.161 | 0.411 |
| 4 | 38892649 | T | C | C | C | 0.275 | 0.94 | 0.956 | 0.642 | 0.07 | 0.097 | 0.825 | 0.161 | 0.411 |
| 4 | 38894579 | C | C | T | C | 0.897 | 0.852 | 0.457 | 0.508 | 0.958 | 0.819 | 0.675 | 0.289 | 0.691 |
| 4 | 38905731 | T | G | G | T | 0.275 | 0.94 | 0.956 | 0.621 | 0.07 | 0.097 | 0.825 | 0.161 | 0.411 |
|  |  | V | III | IV | VII | ITU CHB | ITU JPT | ITU CHS | ITU CDX | ITU KHV | ITU GIH | ITU PJL | ITU BEB | ITU STU |
| 4 | 38760338 | G | T | G | G | 0.658 | 0.94 | 0.047 | 0.871 | 0.782 | 0.262 | 0.871 | 0.379 | 0.171 |
| 4 | 38765046 | A | G | A | G | 0.71 | 0.963 | 0.081 | 0.854 | 0.587 | 0.365 | 0.853 | 0.24 | 0.299 |
| 4 | 38766285 | G | A | G | A | 0.71 | 0.963 | 0.081 | 0.854 | 0.587 | 0.365 | 0.853 | 0.24 | 0.299 |
| 4 | 38767749 | G | A | G | A | 0.951 | 0.993 | 0.778 | 0.148 | 0.059 | 0.979 | 0.153 | 0.93 | 0.382 |
| 4 | 38769018 | A | G | A | G | 0.951 | 0.993 | 0.778 | 0.148 | 0.059 | 0.979 | 0.153 | 0.93 | 0.382 |
| 4 | 38777471 | C | T | C | C | 0.975 | 0.996 | 0.902 | 0.008 | 0.546 | 0.989 | 0.219 | 0.949 | 0.389 |
| 4 | 38778903 | G | A | G | G | 0.975 | 0.996 | 0.902 | 0.008 | 0.546 | 0.989 | 0.219 | 0.949 | 0.389 |
| 4 | 38779512 | C | T | C | T | 0.969 | 0.994 | 0.882 | 0.006 | 0.417 | 0.979 | 0.153 | 0.93 | 0.382 |
| 4 | 38780471 | T | C | T | C | 0.969 | 0.994 | 0.882 | 0.006 | 0.417 | 0.979 | 0.153 | 0.93 | 0.382 |
| 4 | 38783848 | T | C | C | C | 0.984 | 0.995 | 0.952 | 0.534 | 0.733 | 0.918 | 0.146 | 0.762 | 0.287 |
| 4 | 38789675 | G | A | A | A | 0.984 | 0.995 | 0.952 | 0.534 | 0.733 | 0.918 | 0.146 | 0.762 | 0.381 |
| 4 | 38796255 | G | T | T | G | 0.988 | 0.994 | 0.967 | 0.649 | 0.801 | 0.957 | 0.27 | 0.931 | 0.713 |
| 4 | 38802913 | A | G | G | G | 0.99 | 0.998 | 0.974 | 0.561 | 0.824 | 0.953 | 0.157 | 0.873 | 0.381 |
| 4 | 38803063 | A | G | G | G | 0.994 | 0.998 | 0.98 | 0.561 | 0.824 | 0.953 | 0.157 | 0.873 | 0.381 |
| 4 | 38803646 | T | C | C | C | 0.99 | 0.998 | 0.974 | 0.561 | 0.824 | 0.953 | 0.157 | 0.873 | 0.381 |
| 4 | 38803784 | C | T | T | T | 0.99 | 0.998 | 0.974 | 0.561 | 0.824 | 0.953 | 0.157 | 0.873 | 0.381 |
| 4 | 38804405 | C | T | T | T | 0.99 | 0.998 | 0.974 | 0.561 | 0.824 | 0.953 | 0.157 | 0.873 | 0.381 |
| 4 | 38805489 | C | T | T | T | 0.99 | 0.998 | 0.974 | 0.561 | 0.824 | 0.953 | 0.157 | 0.873 | 0.381 |
| 4 | 38805983 | T | C | C | C | 0.99 | 0.998 | 0.974 | 0.561 | 0.841 | 0.953 | 0.163 | 0.873 | 0.381 |
| 4 | 38806019 | A | G | G | G | 0.99 | 0.998 | 0.974 | 0.561 | 0.841 | 0.953 | 0.163 | 0.873 | 0.381 |
| 4 | 38806096 | T | C | C | C | 0.99 | 0.998 | 0.974 | 0.561 | 0.824 | 0.953 | 0.157 | 0.873 | 0.381 |
| 4 | 38806200 | G | T | T | T | 0.99 | 0.998 | 0.974 | 0.561 | 0.824 | 0.953 | 0.157 | 0.873 | 0.381 |
| 4 | 38806827 | G | A | A | A | 0.989 | 0.998 | 0.974 | 0.561 | 0.824 | 0.953 | 0.157 | 0.873 | 0.381 |
| 4 | 38806851 | A | G | G | G | 0.989 | 0.998 | 0.974 | 0.561 | 0.824 | 0.953 | 0.157 | 0.873 | 0.381 |
| 4 | 38807328 | A | G | G | G | 0.988 | 0.998 | 0.968 | 0.534 | 0.784 | 0.918 | 0.146 | 0.762 | 0.287 |
| 4 | 38808045 | A | T | T | T | 0.988 | 0.998 | 0.968 | 0.534 | 0.784 | 0.918 | 0.146 | 0.762 | 0.287 |
| 4 | 38808585 | G | C | C | C | 0.988 | 0.998 | 0.968 | 0.534 | 0.784 | 0.918 | 0.146 | 0.762 | 0.287 |
| 4 | 38808613 | G | A | A | A | 0.988 | 0.997 | 0.968 | 0.534 | 0.784 | 0.918 | 0.146 | 0.762 | 0.287 |
| 4 | 38809924 | G | A | A | A | 0.988 | 0.997 | 0.968 | 0.534 | 0.784 | 0.918 | 0.146 | 0.762 | 0.287 |
| 4 | 38810444 | C | T | T | T | 0.988 | 0.998 | 0.968 | 0.534 | 0.784 | 0.918 | 0.146 | 0.762 | 0.287 |
| 4 | 38810738 | C | G | G | G | 0.988 | 0.998 | 0.968 | 0.534 | 0.784 | 0.918 | 0.146 | 0.762 | 0.287 |
| 4 | 38810769 | C | T | T | T | 0.988 | 0.998 | 0.968 | 0.534 | 0.784 | 0.918 | 0.146 | 0.762 | 0.287 |
| 4 | 38810975 | C | T | T | T | 0.988 | 0.998 | 0.968 | 0.534 | 0.784 | 0.918 | 0.146 | 0.762 | 0.287 |
| 4 | 38811529 | T | A | A | A | 0.988 | 0.998 | 0.968 | 0.534 | 0.784 | 0.918 | 0.146 | 0.762 | 0.287 |
| 4 | 38811668 | G | A | A | A | 0.988 | 0.998 | 0.968 | 0.534 | 0.784 | 0.918 | 0.146 | 0.762 | 0.287 |
| 4 | 38811845 | G | C | C | C | 0.988 | 0.998 | 0.968 | 0.534 | 0.784 | 0.918 | 0.146 | 0.762 | 0.287 |
| 4 | 38812116 | T | A | A | A | 0.988 | 0.998 | 0.968 | 0.534 | 0.784 | 0.918 | 0.146 | 0.762 | 0.287 |
| 4 | 38812537 | A | G | G | G | 0.988 | 0.998 | 0.968 | 0.534 | 0.784 | 0.918 | 0.146 | 0.762 | 0.287 |
| 4 | 38814700 | A | G | G | G | 0.982 | 0.995 | 0.935 | 0.146 | 0.587 | 0.925 | 0.138 | 0.46 | 0.291 |
| 4 | 38827445 | A | A | G | A | 0.849 | 0.448 | 0.493 | 0.963 | 0.81 | 0.654 | 0.086 | 0.716 | 0.111 |
| 4 | 38828383 | A | A | G | A | 0.849 | 0.448 | 0.493 | 0.963 | 0.81 | 0.654 | 0.086 | 0.716 | 0.111 |
| 4 | 38833173 | A | T | T | A | 0.936 | 0.982 | 0.729 | 0.051 | 0.062 | 0.845 | 0.091 | 0.235 | 0.208 |
| 4 | 38835702 | T | C | C | T | 0.936 | 0.984 | 0.729 | 0.051 | 0.062 | 0.845 | 0.091 | 0.235 | 0.208 |
| 4 | 38836071 | G | A | A | G | 0.936 | 0.984 | 0.729 | 0.051 | 0.062 | 0.845 | 0.091 | 0.235 | 0.208 |
| 4 | 38843617 | A | A | T | A | 0.849 | 0.448 | 0.493 | 0.963 | 0.81 | 0.654 | 0.086 | 0.716 | 0.111 |
| 4 | 38843965 | T | T | C | T | 0.849 | 0.448 | 0.493 | 0.963 | 0.81 | 0.654 | 0.086 | 0.716 | 0.111 |
| 4 | 38846188 | T | T | G | G | 0.68 | 0.055 | 0.106 | 0.937 | 0.701 | 0.858 | 0.273 | 0.892 | 0.147 |
| 4 | 38851921 | T | T | G | T | 0.849 | 0.448 | 0.493 | 0.963 | 0.81 | 0.654 | 0.086 | 0.716 | 0.111 |
| 4 | 38853671 | G | G | A | A | 0.68 | 0.055 | 0.106 | 0.937 | 0.701 | 0.858 | 0.273 | 0.892 | 0.147 |
| 4 | 38856365 | T | T | C | T | 0.849 | 0.448 | 0.493 | 0.952 | 0.81 | 0.654 | 0.086 | 0.716 | 0.111 |
| 4 | 38857896 | C | C | A | A | 0.68 | 0.055 | 0.106 | 0.916 | 0.701 | 0.858 | 0.273 | 0.892 | 0.147 |
| 4 | 38871385 | T | C | C | T | 0.927 | 0.976 | 0.634 | 0.028 | 0.061 | 0.73 | 0.021 | 0.289 | 0.182 |
| 4 | 38876997 | G | G | A | G | 0.849 | 0.448 | 0.493 | 0.952 | 0.81 | 0.654 | 0.273 | 0.716 | 0.111 |
| 4 | 38879003 | A | A | C | A | 0.849 | 0.448 | 0.493 | 0.952 | 0.81 | 0.654 | 0.273 | 0.716 | 0.111 |
| 4 | 38881761 | G | A | A | G | 0.912 | 0.98 | 0.535 | 0.028 | 0.133 | 0.718 | 0.157 | 0.366 | 0.189 |
| 4 | 38881766 | G | A | A | G | 0.912 | 0.98 | 0.535 | 0.028 | 0.133 | 0.718 | 0.157 | 0.366 | 0.189 |
| 4 | 38881773 | A | G | G | A | 0.851 | 0.969 | 0.388 | 0.002 | 0.06 | 0.414 | 0.026 | 0.174 | 0.176 |
| 4 | 38886420 | G | A | A | G | 0.912 | 0.932 | 0.523 | 0.003 | 0.037 | 0.44 | 0.02 | 0.17 | 0.189 |
| 4 | 38892649 | T | C | C | C | 0.895 | 0.915 | 0.395 | 0.006 | 0.056 | 0.39 | 0.091 | 0.096 | 0.295 |
| 4 | 38894579 | C | C | T | C | 0.849 | 0.448 | 0.493 | 0.952 | 0.81 | 0.654 | 0.273 | 0.716 | 0.111 |
| 4 | 38905731 | T | G | G | T | 0.912 | 0.932 | 0.395 | 0.003 | 0.037 | 0.44 | 0.02 | 0.17 | 0.189 |

**Supplementary Table 7:**
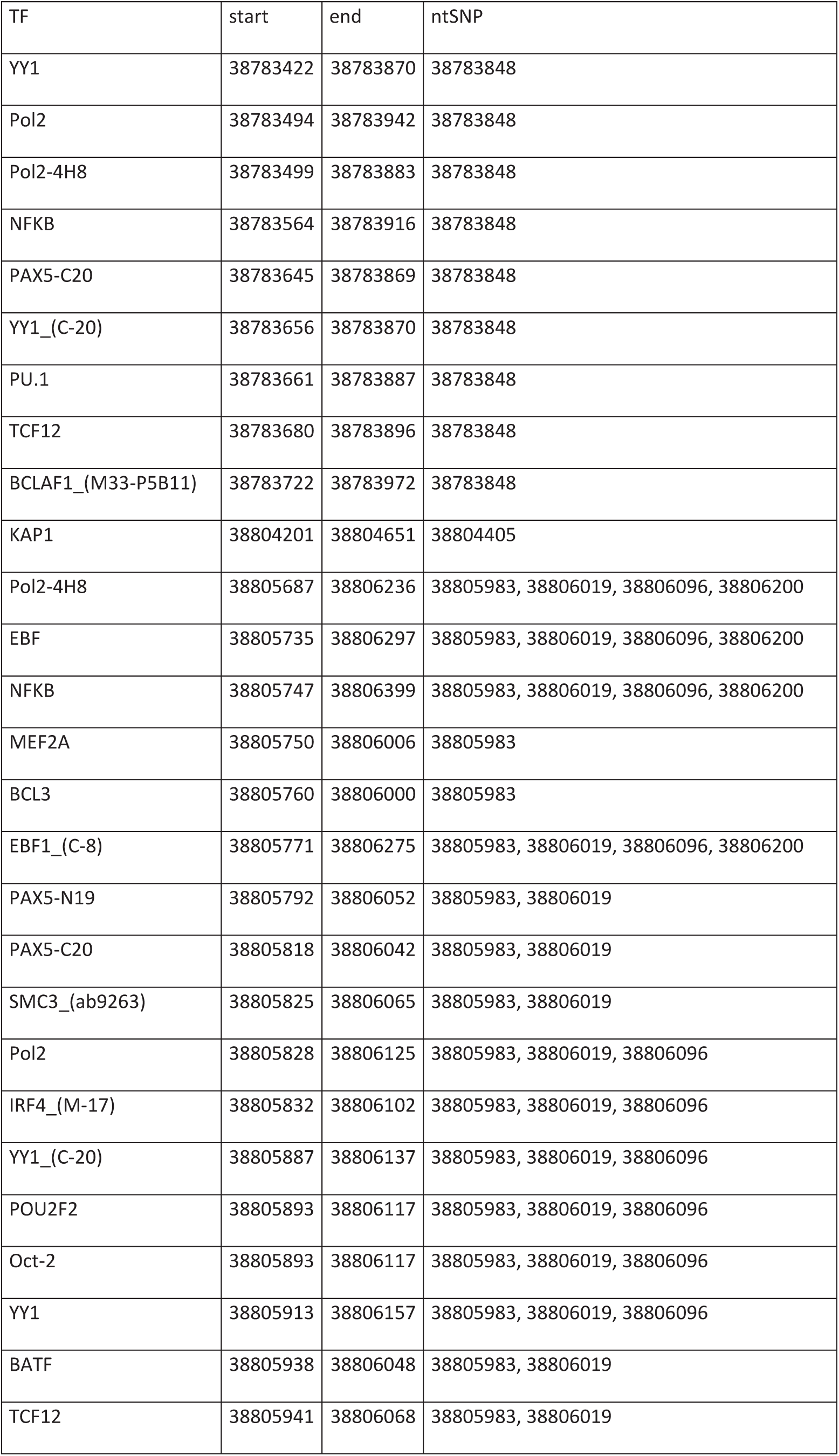
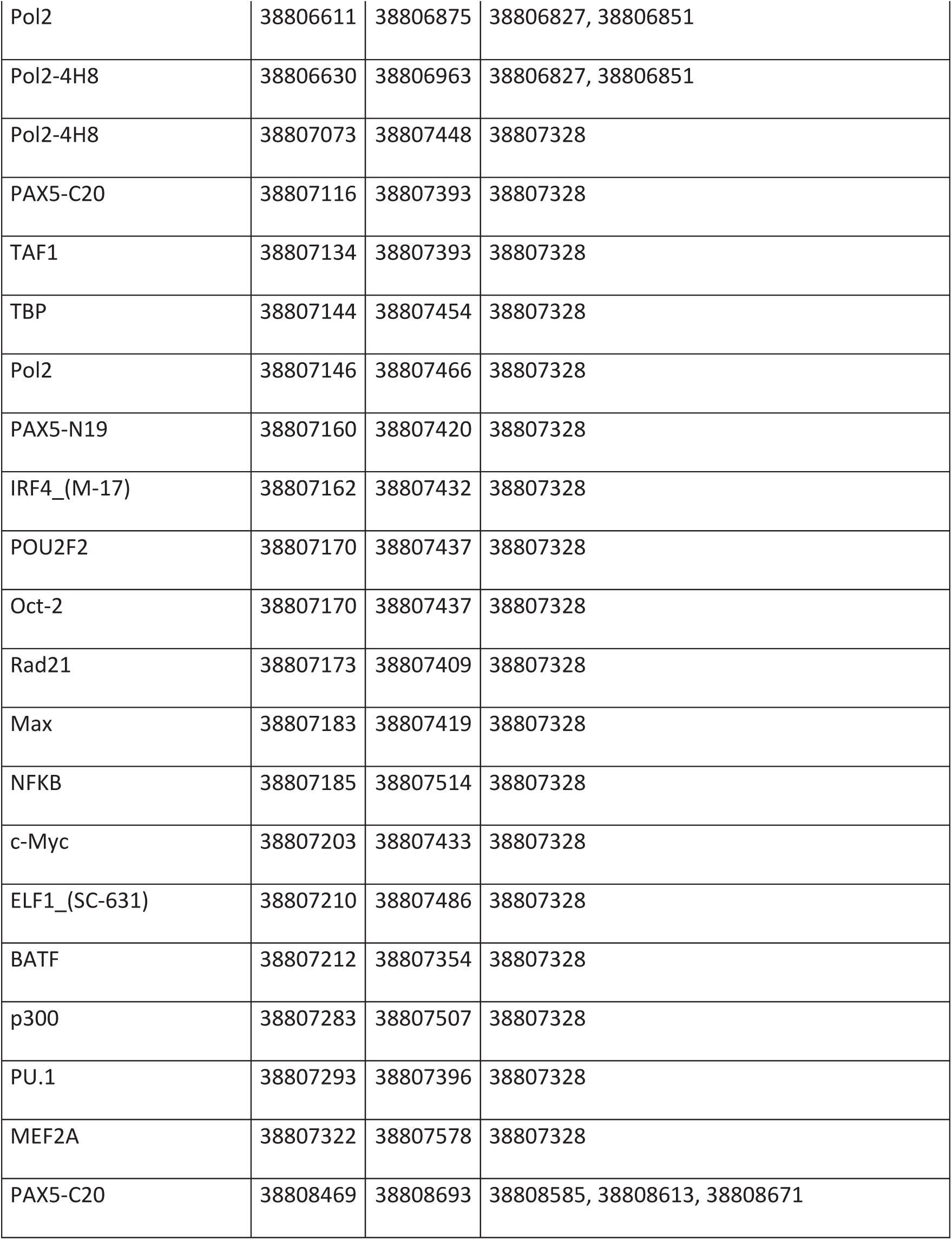
Transcription factor binding sites overlapping archaic-like SNPs

**Supplementary Table 8:**
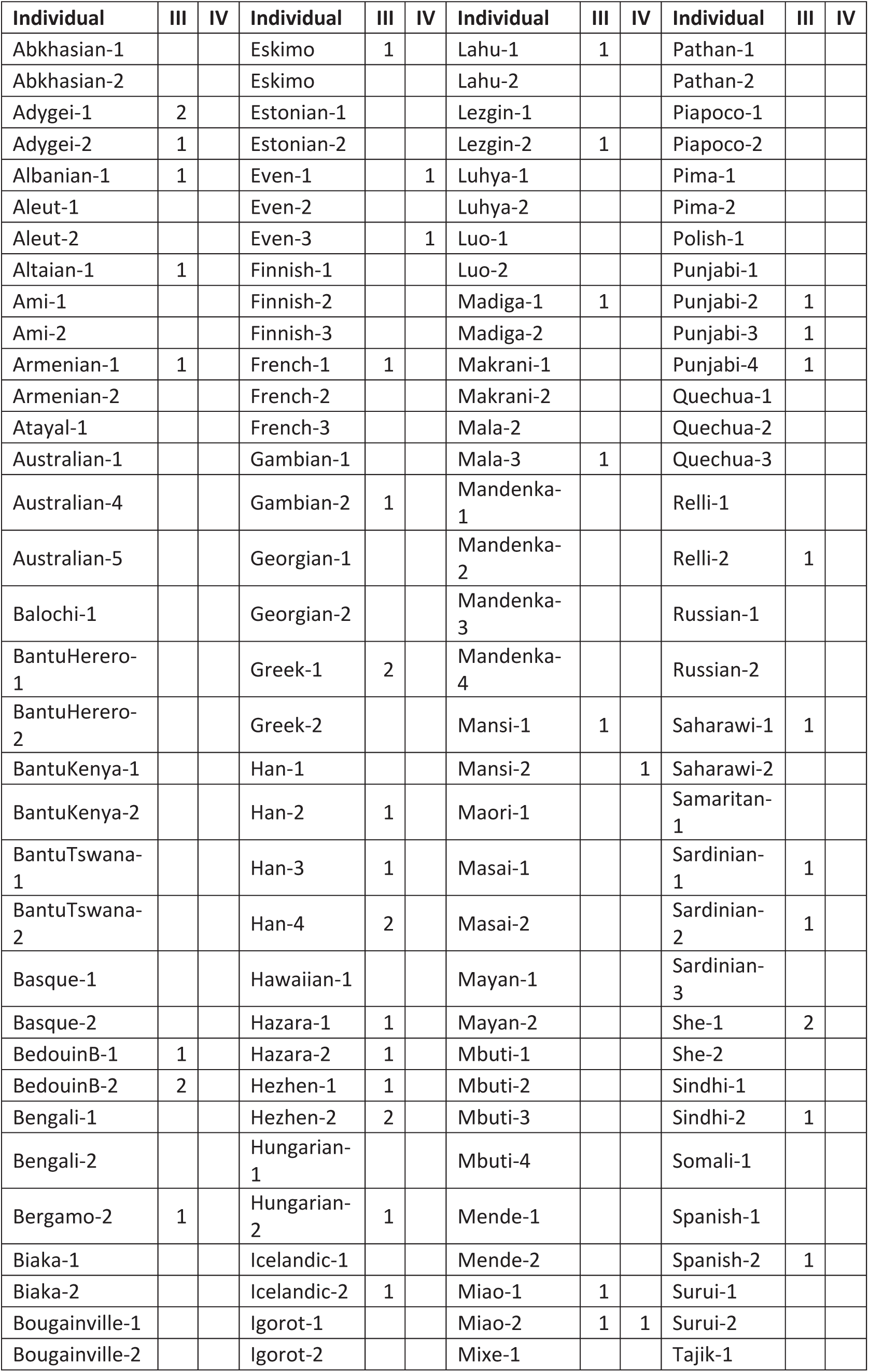
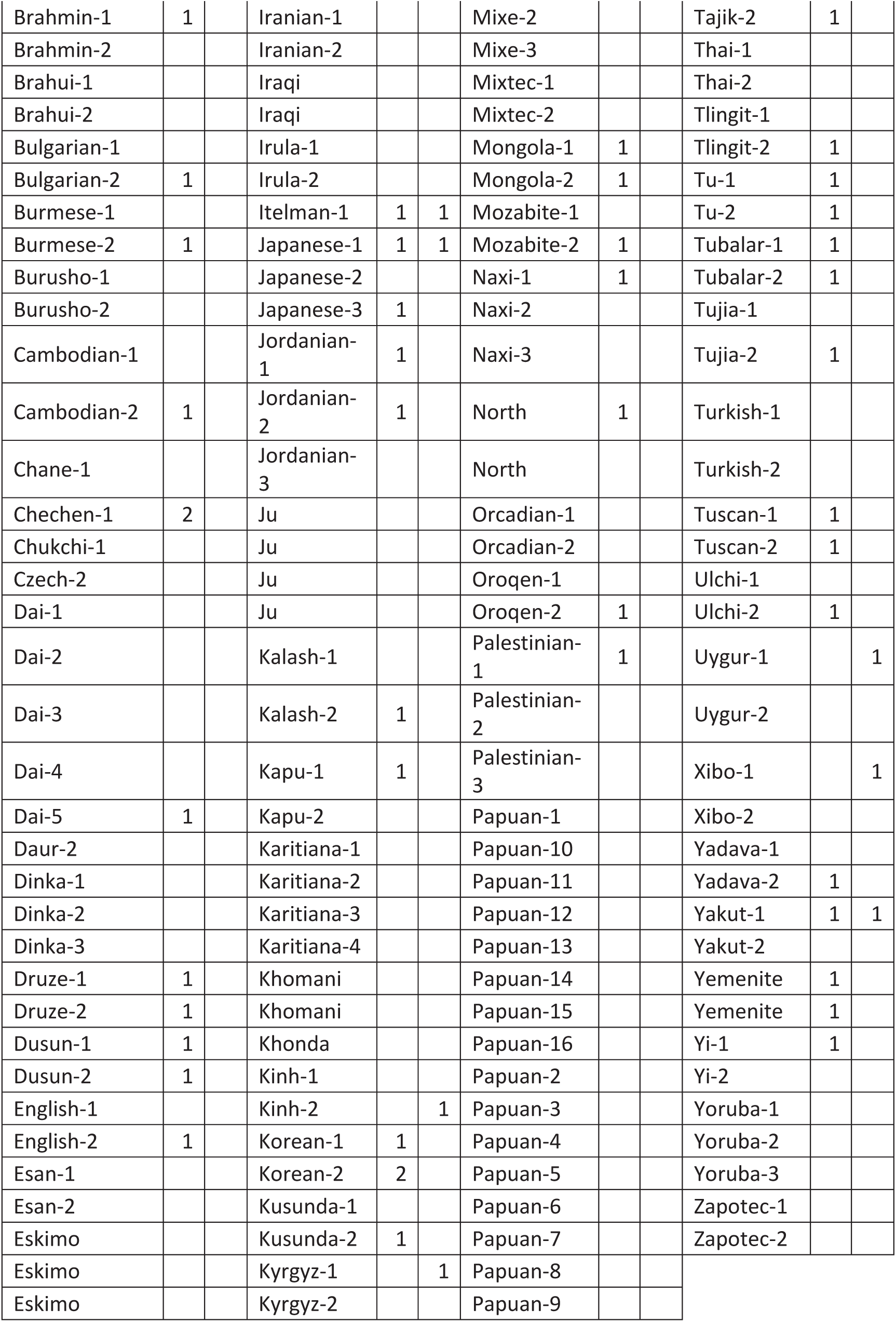
Neanderthal-like core haplotypes in Simons Genome Diversity Panel individuals.

**Supplementary Table 9:** Information on four tag SNPs reported by Barreiro et al. 2009

|  | EUR/ASN | AFR/ASN | AFR/EUR |
| --- | --- | --- | --- |
| 38772827 | - | - | - |
| 38774889<br>rs4129009 | 0.80 | 0.97 | 0.93 |
| 38798648<br>rs5743618 | 1.00 | 0.49 | 1.00 |
| 38799710<br>rs4833095 | 0.97 | 0.97 | 1.00 |
| 38830350<br>rs5743810 | 1.00 | 0.03 | 0.99 |

(b) Majority allele on core haplotypes for tag SNPs
|  | 38774889<br>rs4129009 | 38798648<br>rs5743618 | 38799710<br>rs4833095 | 38830350<br>rs5743810 |
| --- | --- | --- | --- | --- |
| I | T | A | C | G |
| II | T | C | C | G |
| III | C | A | C | G |
| IV | T | A | C | G |
| V | T | A | T | G |
| VI | T | A | C | G |
| VII | T | A | C | G |
| VIII | T | A | C | G |
| IX | T | A | C | G |

(c) Allele frequency on core haplotypes for tag SNPs
|  | 38774889<br>rs4129009 | 38798648<br>rs5743618 | 38799710<br>rs4833095 | 38830350<br>rs5743810 |
| --- | --- | --- | --- | --- |
| I | 100 | 100 | 100 | 100 |
| II | 100 | 0 | 100 | 100 |
| III | 22 | 99 | 91 | 100 |
| IV | 98 | 100 | 97 | 100 |
| V | 99 | 74 | 46 | 85 |
| VI | 98 | 99 | 92 | 100 |
| VII | 100 | 100 | 100 | 100 |
| VIII | 100 | 100 | 100 | 100 |
| IX | 100 | 100 | 100 | 100 |

(d) Allele frequency in modern human populations (1000 genomes phase III) for tag SNPs
|  | 38774889<br>rs4129009 | 38798648<br>rs5743618 | 38799710<br>rs4833095 | 38830350<br>rs5743810 |
| --- | --- | --- | --- | --- |
| CHB | 0.64 | 0.98 | 0.68 | 1.00 |
| JPT | 0.50 | 1.00 | 0.70 | 1.00 |
| CHS | 0.76 | 0.99 | 0.60 | 1.00 |
| CDX | 0.90 | 1.00 | 0.53 | 1.00 |
| KHV | 0.89 | 1.00 | 0.49 | 1.00 |
| GIH | 0.81 | 0.92 | 0.48 | 0.97 |
| PJL | 0.88 | 0.90 | 0.47 | 0.96 |
| BEB | 0.80 | 0.94 | 0.52 | 0.98 |
| STU | 0.86 | 0.95 | 0.54 | 1.00 |
| ITU | 0.90 | 0.93 | 0.55 | 1.00 |
| CEU | 0.81 | 0.23 | 0.21 | 0.52 |
| TSI | 0.69 | 0.47 | 0.40 | 0.66 |
| FIN | 0.91 | 0.17 | 0.15 | 0.58 |
| GBR | 0.84 | 0.23 | 0.20 | 0.54 |
| IBS | 0.72 | 0.50 | 0.43 | 0.64 |
| YRI | 1.00 | 1.00 | 0.90 | 1.00 |
| LWK | 1.00 | 0.99 | 0.93 | 0.99 |
| GWD | 0.99 | 0.98 | 0.86 | 0.99 |
| MSL | 1.00 | 0.99 | 0.93 | 1.00 |
| ESN | 1.00 | 0.98 | 0.89 | 1.00 |
| ASW | 0.93 | 0.86 | 0.77 | 0.92 |
| ACB | 0.98 | 0.91 | 0.84 | 0.96 |
| MXL | 0.92 | 0.82 | 0.49 | 0.85 |
| PUR | 0.77 | 0.59 | 0.45 | 0.78 |
| CLM | 0.86 | 0.63 | 0.45 | 0.74 |
| PEL | 0.94 | 0.88 | 0.51 | 0.92 |

**Supplementary Table 10:** List of the 1000 genomes individuals that are homozygous for core haplotype III

|  |  |  |  |  |
| --- | --- | --- | --- | --- |
| HG00136 | HG01305 | HG04042 | NA18648 | NA19080 |
| HG00141 | HG01378 | HG04146 | NA18944 | NA19084 |
| HG00369 | HG01412 | HG04180 | NA18946 | NA19087 |
| HG00403 | HG01431 | NA12383 | NA18947 | NA19780 |
| HG00419 | HG01522 | NA12751 | NA18949 | NA20510 |
| HG00448 | HG01593 | NA12760 | NA18957 | NA20511 |
| HG00478 | HG01605 | NA12815 | NA18962 | NA20512 |
| HG00531 | HG01606 | NA18525 | NA18967 | NA20517 |
| HG00559 | HG01613 | NA18530 | NA18974 | NA20526 |
| HG00565 | HG01624 | NA18532 | NA18981 | NA20528 |
| HG00590 | HG01678 | NA18539 | NA18982 | NA20540 |
| HG00592 | HG01705 | NA18547 | NA18986 | NA20542 |
| HG00593 | HG01746 | NA18548 | NA18989 | NA20588 |
| HG00614 | HG01756 | NA18550 | NA18990 | NA20756 |
| HG00620 | HG01762 | NA18559 | NA18991 | NA20773 |
| HG00671 | HG01770 | NA18560 | NA18993 | NA20774 |
| HG00698 | HG01775 | NA18566 | NA18994 | NA20783 |
| HG00708 | HG01779 | NA18593 | NA18998 | NA20787 |
| HG00736 | HG01809 | NA18612 | NA19006 | NA20802 |
| HG01028 | HG01847 | NA18615 | NA19055 | NA20804 |
| HG01077 | HG01860 | NA18627 | NA19059 | NA20832 |
| HG01095 | HG02076 | NA18629 | NA19064 | NA20850 |
| HG01097 | HG02775 | NA18633 | NA19065 | NA20854 |
| HG01122 | HG03015 | NA18635 | NA19067 | NA20876 |
| HG01162 | HG03585 | NA18637 | NA19072 | NA20892 |
| HG01168 | HG03594 | NA18638 | NA19074 | NA20893 |
| HG01191 | HG03862 | NA18642 | NA19079 | NA21110 |
| NA21125 | NA21111 |  |  |  |

**Supplementary Table 11:**
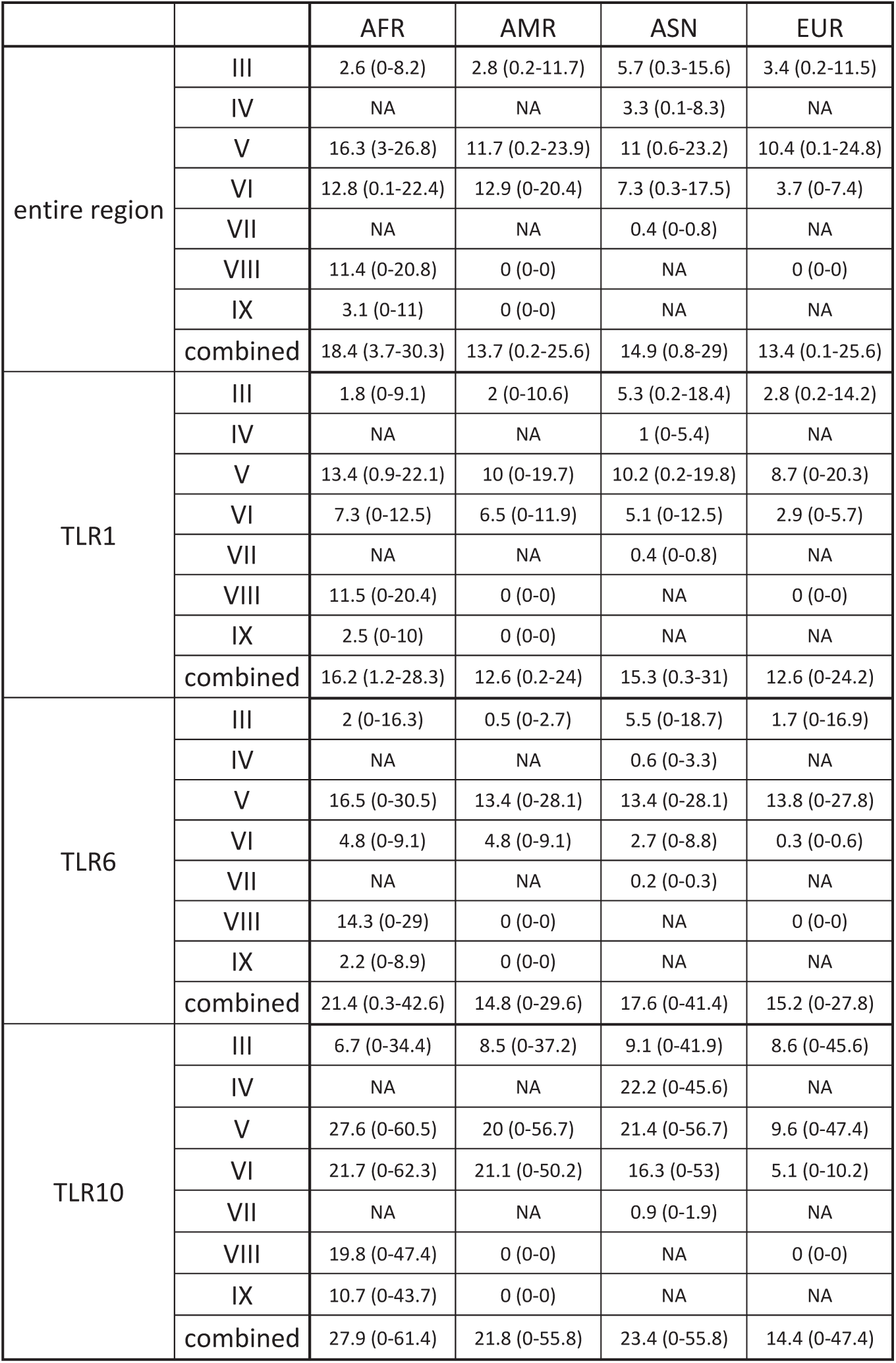
Nucleotide diversity (in 1×10-4 changes per base pair and their 95% quantiles) in three *TLR* genes and the entire *TLR* region across core haplotypes

|  |  | AFR | AMR | ASN | EUR |
| --- | --- | --- | --- | --- | --- |
| entire region | III | 2.6 (0-8.2) | 2.8 (0.2-11.7) | 5.7 (0.3-15.6) | 3.4 (0.2-11.5) |
|  | IV | NA | NA | 3.3 (0.1-8.3) | NA |
|  | V | 16.3 (3-26.8) | 11.7 (0.2-23.9) | 11 (0.6-23.2) | 10.4 (0.1-24.8) |
|  | VI | 12.8 (0.1-22.4) | 12.9 (0-20.4) | 7.3 (0.3-17.5) | 3.7 (0-7.4) |
|  | VII | NA | NA | 0.4 (0-0.8) | NA |
|  | VIII | 11.4 (0-20.8) | 0 (0-0) | NA | 0 (0-0) |
|  | IX | 3.1 (0-11) | 0 (0-0) | NA | NA |
|  | combined | 18.4 (3.7-30.3) | 13.7 (0.2-25.6) | 14.9 (0.8-29) | 13.4 (0.1-25.6) |
| TLR1 | III | 1.8 (0-9.1) | 2 (0-10.6) | 5.3 (0.2-18.4) | 2.8 (0.2-14.2) |
|  | IV | NA | NA | 1 (0-5.4) | NA |
|  | V | 13.4 (0.9-22.1) | 10 (0-19.7) | 10.2 (0.2-19.8) | 8.7 (0-20.3) |
|  | VI | 7.3 (0-12.5) | 6.5 (0-11.9) | 5.1 (0-12.5) | 2.9 (0-5.7) |
|  | VII | NA | NA | 0.4 (0-0.8) | NA |
|  | VIII | 11.5 (0-20.4) | 0 (0-0) | NA | 0 (0-0) |
|  | IX | 2.5 (0-10) | 0 (0-0) | NA | NA |
|  | combined | 16.2 (1.2-28.3) | 12.6 (0.2-24) | 15.3 (0.3-31) | 12.6 (0-24.2) |
| TLR6 | III | 2 (0-16.3) | 0.5 (0-2.7) | 5.5 (0-18.7) | 1.7 (0-16.9) |
|  | IV | NA | NA | 0.6 (0-3.3) | NA |
|  | V | 16.5 (0-30.5) | 13.4 (0-28.1) | 13.4 (0-28.1) | 13.8 (0-27.8) |
|  | VI | 4.8 (0-9.1) | 4.8 (0-9.1) | 2.7 (0-8.8) | 0.3 (0-0.6) |
|  | VII | NA | NA | 0.2 (0-0.3) | NA |
|  | VIII | 14.3 (0-29) | 0 (0-0) | NA | 0 (0-0) |
|  | IX | 2.2 (0-8.9) | 0 (0-0) | NA | NA |
|  | combined | 21.4 (0.3-42.6) | 14.8 (0-29.6) | 17.6 (0-41.4) | 15.2 (0-27.8) |
| TLR10 | III | 6.7 (0-34.4) | 8.5 (0-37.2) | 9.1 (0-41.9) | 8.6 (0-45.6) |
|  | IV | NA | NA | 22.2 (0-45.6) | NA |
|  | V | 27.6 (0-60.5) | 20 (0-56.7) | 21.4 (0-56.7) | 9.6 (0-47.4) |
|  | VI | 21.7 (0-62.3) | 21.1 (0-50.2) | 16.3 (0-53) | 5.1 (0-10.2) |
|  | VII | NA | NA | 0.9 (0-1.9) | NA |
|  | VIII | 19.8 (0-47.4) | 0 (0-0) | NA | 0 (0-0) |
|  | IX | 10.7 (0-43.7) | 0 (0-0) | NA | NA |
|  | combined | 27.9 (0-61.4) | 21.8 (0-55.8) | 23.4 (0-55.8) | 14.4 (0-47.4) |

**Supplementary Table 12:** Probability of archaic haplotype block from shared ancestral lineage using different mutation rates

|  |  |  |  |
| --- | --- | --- | --- |
| Branch lengths since divergence (years) | Human-Neandertal/Denisova | 275000 | 550000 |
|  | Neandertal-Denisova | 190000 | 381000 |
| Recombination rate | per base pair per year | $1 \times 10^{-9}$ | $0.5 \times 10^{-9}$ |
| Age of the Neandertal | Years | 68000 | 136000 |
| Age of the Denisovan |  | 54000 | 108000 |
| Expected length (bp) | Neandertal | 4198.15 | 2096.44 |
|  | Denisova | 4055.15 | 2025.11 |
| Probability of observing a block of at least 143000 bp | Neandertal | $< 2.2 \times 10^{-16}$ | $< 2.2 \times 10^{-16}$ |
| | Denisova | $< 2.2 \times 10^{-16}$ | $< 2.2 \times 10^{-16}$ |

